# A multimodal 3D neuro-microphysiological system with neurite-trapping microelectrodes

**DOI:** 10.1101/2021.04.16.436793

**Authors:** Beatriz Molina-Martínez, Laura-Victoria Jentsch, Fulya Ersoy, Matthijs van der Moolen, Stella Donato, Torbjørn V. Ness, Peter Heutink, Peter D. Jones, Paolo Cesare

**Affiliations:** NMI Natural and Medical Sciences Institute at the University of Tübingen, 72770 Reutlingen, Germany; German Center for Neurodegenerative Diseases (DZNE) & Hertie Institute for Clinical Brain Research, 72076 Tübingen, Germany; Faculty of Science and Technology, Norwegian University of Life Sciences, 1432 Ås, Norway

**Author notes:** Equal contributions (Beatriz Molina-Martínez, Laura-Victoria Jentsch). CICbiomaGUNE, Paseo de Miramón 182, 20014 Donostia-San Sebastián, Spain (Beatriz Molina-Martínez).

## Abstract

Three-dimensional cell technologies as pre-clinical models are emerging tools for mimicking the structural and functional complexity of the nervous system. The accurate exploration of phenotypes in engineered 3D neuronal cultures, however, demands morphological, molecular and especially functional measurements. Particularly crucial is measurement of electrical activity of individual neurons with millisecond resolution. Current techniques rely on customized electrophysiological recording set-ups, characterized by limited throughput and poor integration with other readout modalities. Here we describe a novel approach, using multiwell glass microfluidic microelectrode arrays, allowing non-invasive electrical recording from engineered 3D neural tissues. We demonstrate parallelized studies with reference compounds, calcium imaging and optogenetic stimulation. Additionally, we show how microplate compatibility allows automated handling and high-content analysis of human induced pluripotent stem cell–derived neurons. This microphysiological platform opens up new avenues for high-throughput studies on the functional, morphological and molecular details of neurological diseases and their potential treatment by therapeutic compounds.

## Introduction

Despite recent decades’ impressive progress in treating diseases of many organ systems, brain disorders remain mostly unsolved. Failures of prospective drugs in clinical trials reflect the challenging complexity of the nervous system but may also stem from partially inadequate preclinical models. Well-established two-dimensional (2D) cultures of adherent cells reliably mimic certain aspects of neuronal dysfunctions with high throughput^1^. Yet their results are not leading to effective treatments. Engineered three-dimensional (3D) culture models that better mimic neural tissue present one path towards levelling the translational gap from in vitro to in vivo studies^2,3^.

To generate in vitro models that better recapitulate important features of neuronal networks, recent approaches combine microfluidics (MF) and three-dimensional (3D) culture techniques^4,5^. Gene expression in 3D cultured neurons matches in vivo patterns more closely than monolayer cultures^6–8^, possibly due to 3D interactions and a more natural spatial organization of synaptic connections^9,10^. These 3D models vary in the distribution of the neurons^11^ and the integration or absence of a scaffold^12^. Cerebral organoids, for instance, are 3D self-arranged stem cell-derived neuronal tissues^11,13^ that have attracted attention as they partially recapitulate brain development. However, their compact 3D multicellular architecture challenges functional readout, with most approaches relying on either optical techniques^14^ or planar microelectrodes that record from superficial neurons^15,16^. Alternatively, 3D cultures can distribute dissociated, non-adherent neurons in a hydrogel or scaffold. This more adaptable culture approach mimics a tissue-like environment, allows control over cell type and density, and enables analysis of the 3D network by combining microfluidic designs with microscopy-based readouts, such as calcium imaging^17^. However, direct readout of electrical activity – the most relevant measure of neuronal function – remains challenging. Complex devices such as multi-shank 3D neural probes allow readout of 3D neural network activity^18^, but are poorly suited for the demands of high throughput automated culture. To fill this gap, we report here a neuro-microphysiological system (nMPS), based on microelectrode arrays (MEA) and glass microfluidics^19^, capable of electrophysiological readout of 3D neuronal circuits in a microplate-compatible format.

A core innovation is the addition of insulating caps on substrate-integrated microelectrodes (capped microelectrodes, CME), inspired by tunnels which enable neurite recordings from adherent neurons^20,21^. Our system enables 3D cell co-culture in a hydrogel scaffold with unguided extension of neurites into the CME. We investigated this new capability to use a 2D MEA to record the electrical activity of 3D neuronal circuits with calcium imaging and optogenetic stimulation. We validated the nMPS as a screening platform with the neurotoxins picrotoxin and tetrodotoxin; recorded data showed excellent sensitivity and experimental reproducibility. Electrophysiological read-out identified effects of rotenone, a neurotoxic insecticide that induces a Parkinson’s disease (PD) phenotype in animal models^22^, with superior sensitivity and at earlier time-points than morphological or metabolic in vitro assays. The nMPS enables the acquisition of morphological data from subcellular structures up to full 3D networks, and we have demonstrated how its microplate-compatibility enables automated handling for improved throughput of cell culture and high-content imaging. We further evaluated the system with neurons derived from human induced pluripotent stem cells (hiPSC), supporting its use for modelling human physiology and disease. Enabling the engineering of cell type, density and 3D organization of neuronal networks in a high-throughput platform for functional and structural analysis represents a major step towards more predictive preclinical models to facilitate the translation of therapies for human neurodegenerative disorders.

## Results

### Designing a multi-well 3D nMPS

Strict requirements for multimodal readout guided the development of the nMPS. The microfluidic device needed to support simultaneous morphological and functional assessment, have multiple independent wells, and enable long-term culture lasting at least several weeks. Functionality of the CME required precise microfabrication, which demanded materials with sufficient thermal stability (170 °C), compatibility with organic solvents, and biocompatibility. Therefore, glass and quartz were preferred over thermoplastics or siloxanes. The material constraints provided the benefit of reusability. We designed multiwell microfluidic components with microscale 3D structures, which were produced by selective laser-induced etching (SLE) of quartz (Fig. 1a). Exploring multiple chip architectures allowed convergence on a design that achieves simple cell handling in 18 independent wells (Fig. 1b). Each well (Fig. 1c) has a gel layer (GL) in which dissociated brain cells are dispersed and whose shape is defined by surface tension. After hydrogel polymerization, a liquid layer (LL) of culture medium covers the GL for exchange of oxygen and nutrients. The system achieved long-term viability of 3D cultures with static methods as established in 2D cultures, without any external perfusion systems commonly used in organ-on-a-chip devices^23^. During maturation of the culture, a complex 3D multi-cellular architecture takes shape in each well. This was visualized by live imaging (Fig. 1d–e, supplementary videos 1 and 2) and immunocytochemical methods (Fig 1f–h, supplementary videos 3, 4 and 5) adapted to label cells in the ∼1-mm-thick hydrogel. Confocal imaging revealed healthy cells homogeneously dispersed in 3D in each well. Neurons extended neurite arborizations in all directions (Fig. 1d) with visible dendritic spines (Fig. 1e). Non-neuronal cells (astrocytes and microglia) developed natural morphologies more closely than 2D cultures (Fig. 1f). Axons and dendrites (Fig. 1g) formed a neuronal network with synaptic structures (Fig. 1h) dispersed in the three-dimensional space.

**Figure 1:**
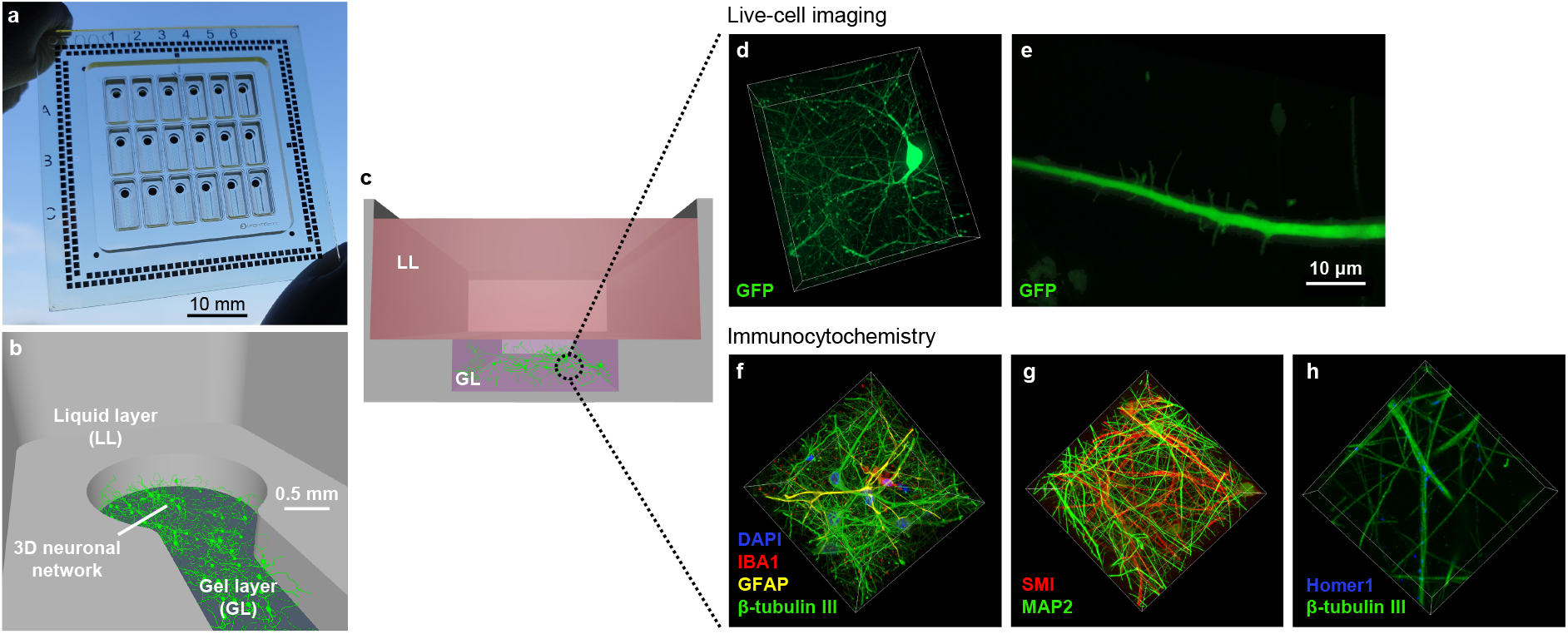
Culturing 3D brain cells in the nMPS. **a**, nMPS with 18 independent wells bonded to the MEA substrate. **b**, Neurons are embedded in a 3D scaffold within the gel layer (GL). Culturing medium is added to the liquid layer (LL). **c**, Cross-section of a single well. **d**, Live-cell imaging of an eGFP-transduced 3D neuron (dimensions 134×151×45 μm) and **e**, a neurite growing dendritic spines. Scale bar 10 µm. **f**, Immunocytochemistry of neurons (β-tubulin III) co-cultured in 3D with astrocytes (GFAP) and microglia (IBA1), with their nuclei stained by DAPI. Dimensions 112×112×109 μm. **g**, Double immunolabeling of axons (SMI) and dendrites (MAP2) forming a complex neuronal network. Dimensions 112×112×73 μm. **h**, 3D immunostaining of postsynaptic density-related marker (Homer 1) and neurite-related marker (β-tubulin III). Dimensions 47×47×11 μm.

### Measuring 3D neuronal activity with CME

Recreating and understanding 3D cell-to-cell interactions and morphology in vitro opens new opportunities to model brain disorders. However, electrical activity of neural circuits forms the basis of cognitive, sensory, and motor functions. Current electrophysiological techniques, either based on glass micropipettes^24^ or on substrate-integrated microelectrodes^25^, are poorly suited for measuring the activity of neurons cultured in 3D. These approaches may be damaging to the recorded neurons, or fail to resolve extracellular signals of cells dispersed in 3D (Fig. 2a, open electrode). To fill this gap, we implemented a microelectrode-based method specifically for recording from non-adherent 3D neurons. Our method introduces the concept of capped microelectrodes (CME), in which conventional substrate-integrated microelectrodes (30 µm diameter) are insulated with a thin (20 µm) cap containing tunnels (5 µm wide and 250 µm long). Neurons cultured in a 3D hydrogel extend neurites in all directions, including into the CME tunnels (Fig. 2a). When neurons fire action potentials, the electrical signals travel along neurites and into the caps. Within a cap, the restricted spread of transmembrane currents generates extracellular action potentials of up to hundreds of microvolts. In contrast, the small extracellular signal (∼5 µV) generated near a neurite cannot be distinguished from recording noise by an adjacent open microelectrode. Fig. 2b demonstrates our concept by simulation of the extracellular potential as an action potential travels along an axon near open and capped microelectrodes (see also Supplementary video 6). Membrane currents were based on an unmyelinated model^26^ and extracellular potentials were simulated using LFPy 2.0^27^ and NEURON^28^ with hydrogel conductivity of 1.46 S/m. Under such conditions, extracellular amplitudes exceeding a threshold of 10 µV are reached only within 15 µm of the soma. As the average cell spacing with a 3D density of 1000–3000 cells per µl is 70–100 µm, standard open electrodes struggle to resolve activity in 3D cultures.

**Figure 2:**
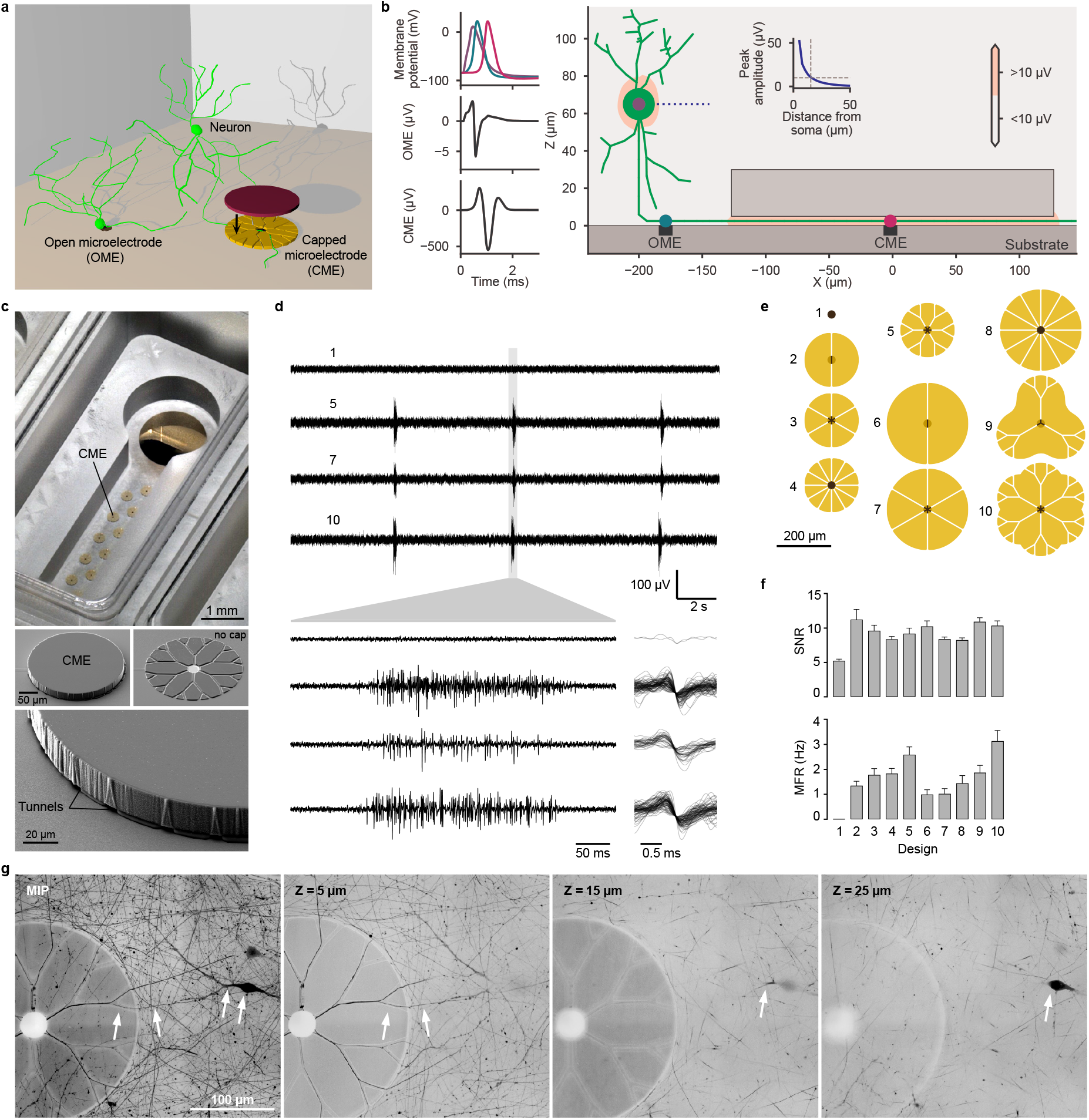
A novel approach to recording electrical activity from 3D neuronal networks. **a**, Schematic drawing of possible configurations to record electrical activity with substrate-integrated microelectrodes from neurons growing in a 3D environment. Neuronal soma can be located nearby an open microelectrode (OME) at the bottom or more dispersed in the 3D volume, projecting its neurites through the tunnels towards the CME. **b**, Comparison of different recording simulations. Graphs indicate membrane potentials at the soma (purple), proximal segment of the axon (green), and distal segment of the axon (pink), as well as extracellular potentials at the OME and the CME. The inset graph shows the peak extracellular potential amplitude as a function of the distance from the soma; amplitudes above noise level (10 µV) are only reached within a distance of 15 µm. **c**, CMEs are placed along the gel layer of a well. Scanning electron microscopy (SEM) reveals the 3D structure of CME and tunnels. **d**, Comparison between the electrical activity measured by various CME designs (5, 7, 10) and an open electrode (1). **e**, Overview of an open electrode (1) and capped electrode designs (2-10). **f**, The average signal-to-noise ratio (SNR) and the mean firing rate (MFR) captured by capped electrodes. Test recordings consist of 3 min measuring the spontaneous electrical activity of primary hippocampal neurons after 8 DIV (N = 3). **g**, A non-adherent neuron extends its neurites into the tunnels of a CME, showing a maximum intensity projection and three confocal images at *Z* heights of 5, 15 and 25 µm. Images are inverted for clarity.

We tested how different cap geometries (Fig. 2e) affect the signal-to-noise ratio (SNR) and the mean firing rate (MFR). The designs varied in their number of tunnel entrances (2 to 24) and diameter (150 to 300 µm) and were compared to open electrodes (Fig. 2e, design 1). Fig. 2d shows representative recordings from 3D-cultured primary mouse hippocampal neurons cultured using designs 1, 5, 7 and 10. Different CME designs resulted in mean SNR of 8.2 to 11.2 and MFR of 0.99 to 3.12 Hz (Fig. 2f), depending on the ease of access for neurites (number of tunnel entrances) and electrical resistance of the tunnels (number and length of tunnels). We detected negligible action potentials at open electrodes (design 1) over the entire period (MFR = 0.01 ± 0.002 Hz), confirming their unsuitability for recording from neurons dispersed in a 3D hydrogel. Confocal microscopy of live cells after fluorescent labelling (AAV-hSyn-EGFP) confirmed that neurites growing through tunnels in fact come from cell bodies at different heights in the 3D neuronal network (Fig. 2g).

### Uniform collection of electrophysiological data across synaptic circuits

A 3D multicellular architecture does not necessarily imply that individual neurons form a functional network. Synchronized action potentials recorded by CMEs (Fig. 2d) suggest that neurons are synaptically connected. However, additional information about the source of such activity is required to establish a spatial correlation between the recorded signal and excitability within the 3D network. Considering the electrodes’ locations (integrated into the glass substrate), it could be argued that they may preferentially record from neurons near the bottom of the 3D culture. To shed light on this, we performed calcium imaging simultaneously with electrical recordings (Fig. 3a and 3b) to identify active neurons within the 3D network. With calcium imaging, activity was measured at a focal plane at least 500 µm above the electrodes. Cells were recorded under basal conditions and following application of picrotoxin (PTX) and tetrodotoxin (TTX), which increase or reduce neuronal network activity, respectively. Network bursts (NB in Fig. 3a) recorded by CME occurred concurrently with large intracellular calcium transients triggered by synchronous activity in neurons (Fig. 3b), which was more evident with PTX-evoked seizure-like activity. Overall, this demonstrates the CME recordings represent activity of the entire volume of the 3D neuronal network.

**Figure 3:**
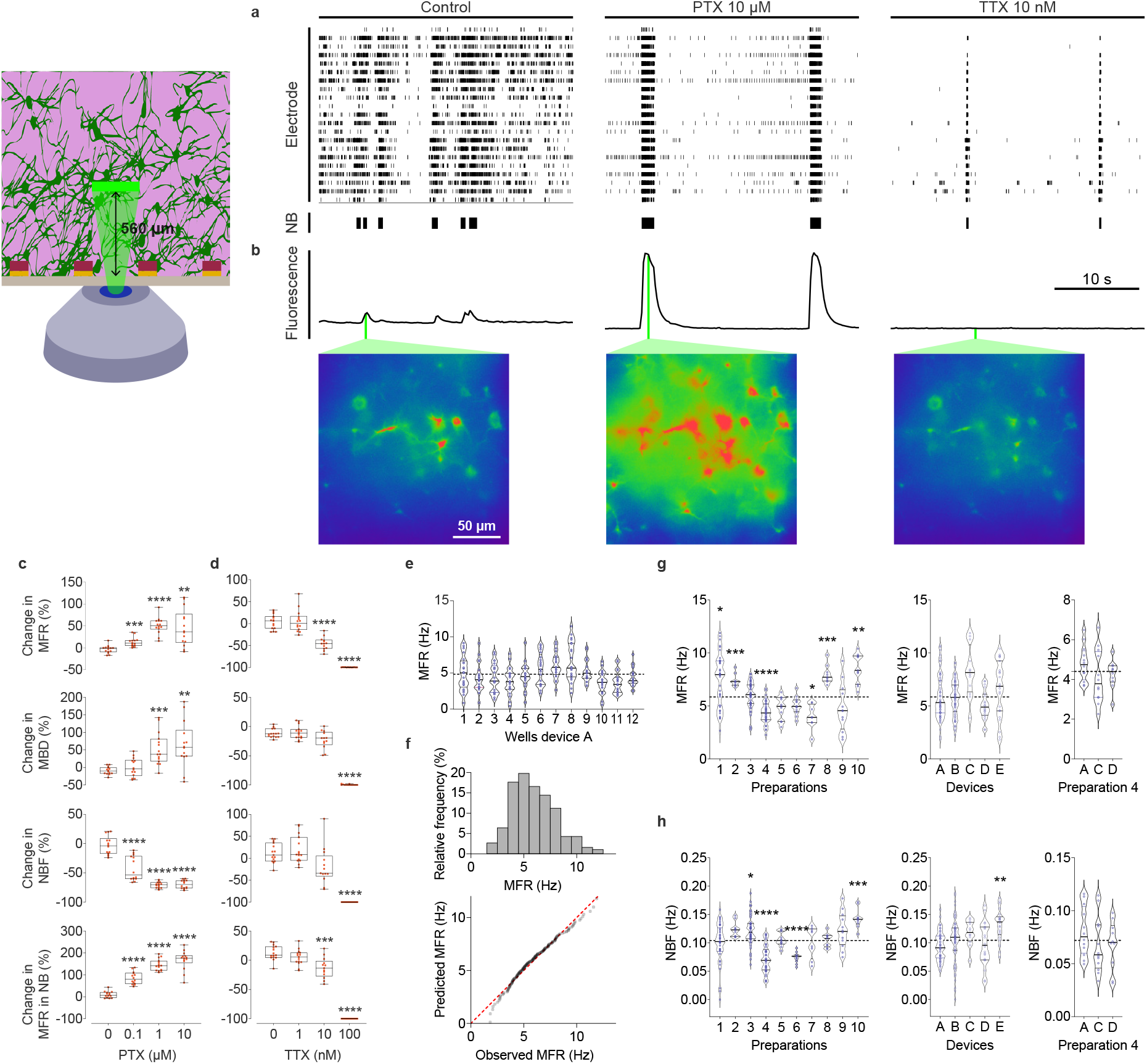
Functional validation of the nMPS. **a**, Raster plots of control sample (left), followed by treatment with 10 µM PTX (middle) and 10 nM TTX (right). Electrical activity of 3D neuronal networks was captured from single wells. Network bursts are illustrated below as black bars. **b**, Fluorescence traces of simultaneous Ca^2+^-imaging of GCaMP6f-transduced neurons at a focal plane of 560 µm. Color map images indicate the calcium intensity response in a ROI of 200×200 µm. The strongest response is shown in red and the weakest response in blue. Scale bar, 50 µm. **c, d**, Box plots representing the change in mean firing rate (MFR), mean burst duration (MBD), network burst frequency (NBF), and mean firing rate in network bursts (MFR in NB) for PTX **(c)** and TTX **(d)**. The line within each box represents the median. The whiskers display the maximum and minimum values, and the interquartile range is set at 90%. This comparative experiment was performed in triplicate with multiple devices. Each red dot represents the value of a single well. **e**, Intervariability of wells in one 12-well device is displayed in a violin plot illustrating the MFR of the spontaneously active 3D neuronal networks. Green dots indicate the single value of each covered electrode. **f**, Collected MFR data of spontaneous activity recorded under the same conditions is represented in a frequency histogram and QQ plot to gauge the degree of normality of the data distribution (N = 187 wells). **g, h**, Representation of the MFR (g) and NBF (h) in violin plots compare the activity recorded in wells grouped by preparation days (left), devices (middle), and devices employed with cells from the same preparation (right). Black dashed lines in plots indicate the mean value. Blue dots represent the value of single wells. Significance between the mean value and the mean of each group is displayed as asterisks: *p<0.05; **p<0.01; ***p<0.001; ****p<0.0001.

### Validating reproducibility of nMPS data by reference compounds

Both electrical and image-based readouts provide crucial information about neuronal activity. However, in comparison to calcium imaging, recordings by CME offer the advantage of direct, non-invasive, and highly sensitive measurements of the electrical activity across an entire 3D neuronal network. We validated our electrophysiological readout by quantifying the response to different concentrations of PTX and TTX. Specifically, we quantified the mean firing rate (MFR), mean burst duration (MBD), network burst frequency (NBF), and mean firing rate in network bursts (MFR in NB) (Figures 3c, d), among others (Supplementary figure 2). Low concentrations (0.1 µM) of PTX significantly affected the network activity by increasing MFR and MFR in NB. Additionally, PTX decreased NBF and MBD, although the latter showed a significant effect only at concentrations of 1 and 10 µM. TTX significantly reduced the neuronal activity by reducing all MBD, MFR, NBF, and MFR in NB, but only at higher concentrations.

We examined experimental reproducibility using the nMPS by comparing spontaneous activity measured under control conditions across different wells, devices and preparations. Low intrinsic variability of MFR (Fig. 3e) and other parameters (supplementary figure 3) was confirmed by evaluating the inter-well variability. Although firing rates recorded from different electrodes in the same well were variable, the mean and median firing rates were similar (3.7 to 6.5 Hz) when compared between independent wells in each device. This indicated that 3D neuronal cultures seeded at the same time in a given device generate comparable data with low variability. We confirmed that MFR and other electrophysiological parameters followed a normal distribution, with a slight leftwards skew, as seen in a histogram and quantile-quantile plot (Fig. 3f and supplementary figure 4) and characteristic of mature neuronal networks^29,30^.

In comparison to the variability for multiple wells of a single preparation, we observed higher variability of MFR and other measures (supplementary figure 5) in independent biological preparations. Our analysis indicates that this variability does not arise from technical issues related to the different nMPS devices (Fig. 3g, supplementary figures 6 and 7). We observed similar patterns of variability for NBF (Fig. 3h). Overall, we found that the dominant source of variability in spontaneous activity originated from the cell preparation. This variability may result from both biological differences in our animal-derived primary cultures and technical differences during cell isolation and seeding.

### Modulating excitability in 3D neuronal networks by optogenetic stimulation

We next investigated the use of optogenetic stimulation with the nMPS. Neuronal cultures were transduced by AAV particles expressing channelrhodopsin-2 (hChR2), a cation-selective ion channel which causes rapid and reversible depolarization when activated by short (millisecond) pulses of blue light (470 nm). The microscope objective was focused at different heights in the 3D culture, and a defined region between electrodes was targeted by patterned illumination with a digital micromirror device (Fig. 4a). The electrical activity of the 3D network was recorded during sequences of light pulses (forty 5-ms pulses at 40 Hz every 10 seconds). This optical activation of neurons in the illuminated area triggered activation of action potentials and synchronized network bursts across the whole well as measured by individual electrodes (Fig. 4b). This confirms that the CME, despite being integrated at the bottom of the nMPS, can represent the electrical activity of the whole 3D neuronal network. Additionally, it proves that neurons cultured in 3D in the nMPS form functional synapses, express ion channels and sustain different excitability patterns (Fig. 4c). To explore this capacity at higher throughput, all hChR2-expressing neurons of a whole device were stimulated by illuminating all wells simultaneously (Fig. 4d). Wells exhibited independent spontaneous activity, as expected. Then, each stimulation sequence generated simultaneous network bursts across all wells (Fig. 4e). Despite different basal activity under resting conditions, neurons in all wells reacted similarly when stimulated. These results further demonstrate the reproducibility of the nMPS for establishing and recording from 3D neuronal networks. We anticipate that the nMPS, in combination with high-frequency optical stimulation, could be used to explore mechanisms of synaptic plasticity (long-term potentiation and depression) in 3D neuronal circuits in vitro with a throughput compatible with drug discovery.

**Figure 4:**
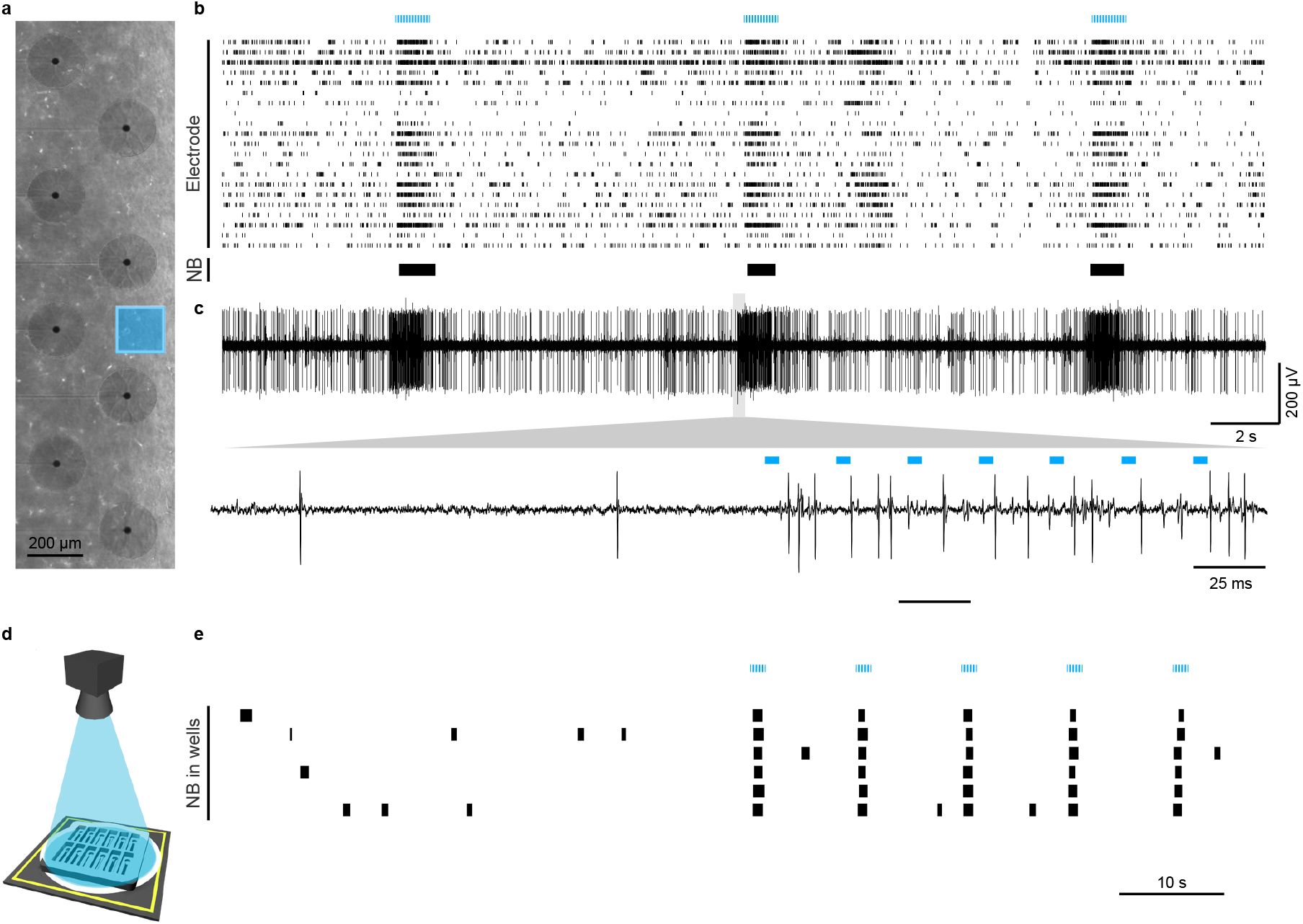
Optical stimulation of 3D neuronal networks. **a**, hChR2-expressing neurons were illuminated with blue light (λ = 470 nm) within a defined area (200×200 µm) at a height of 100 µm from the bottom of the 3D scaffold. **b**, The electrical activity of the whole neuronal network was continuously monitored by the integrated electrodes, while light pulses (5 ms) were repeatedly applied for 1 s (at 40 Hz) every 10 seconds. Neuronal activity captured by the 21 integrated electrodes of one single well (12-well nMPS) is shown as a raster plot. Blue lines indicate the time window of the optical stimulation and bold black lines represent network bursts. **c**, Trace plots representing the activity recorded by one CME at different time scales. **d**, Illustration of the multi-well optical stimulation approach employing a fiber optic. **e**, Raster plot of network bursts evoked in six wells by simultaneous light stimulation (5 ms pulses, 1 s at 40 Hz, repeated every 10 s).

### Integrating electrophysiological and morphological readouts into a 3D neurotoxicity assay

The nMPS consolidates electrophysiological readout and morphological analysis in a multiwell microplate-compatible design. This represent a true novelty for 3D neuronal networks, as current 3D in vitro neuronal assays only rely on morphological^17,31^ or low-throughput, invasive electrophysiological readouts^18,32^. We demonstrate the benefits of combining functional and structural assays for in vitro investigation of neurodegenerative disorders by examining the effects of rotenone on mature (more than 7 DIV) 3D neuronal cultures. Rotenone is a pesticide that inhibits the mitochondrial complex I and causes alpha-synuclein accumulation^33^. Its application in neuronal cultures models alterations associated with Parkinson’s disease (PD), with effects on neurite outgrowth and metabolic stress^22,34^. Morphology-based biomarkers cannot measure subtle changes in excitability and synaptic transmission. Failing to predict these neurologically relevant effects reduces the translational value of such in vitro studies. To fill this gap, we used the nMPS to assess excitability in 3D neuronal cultures exposed to rotenone (0.05–5 µM). Surprisingly, even the lowest dose decreased neuronal activity (Fig. 5a). This effect was confirmed by significant decreases in MFR, MBD, NBF and MFR in NB (Fig. 5b) (the large deviation of MFR in NB at higher concentrations results from the diminished occurrence of NB). In parallel, we studied the morphology of rotenone-treated 3D neuronal cultures by live-cell imaging at different time-points (20 min, 6 h, and 24 h) (Fig. 5c). Although changes in electrical activity were detected just 10 min after the application of 0.05 µM rotenone, morphological alterations only appeared after 6 h and 24 h at a concentration of 5 µM and 0.5 µM, respectively. This confirms that electrophysiological assessment of 3D neuronal networks can supplement morphological studies and provide a more sensitive approach to study the effects of neuroactive and toxic compounds, and thereby enhance predictability of in vitro methods.

**Figure 5:**
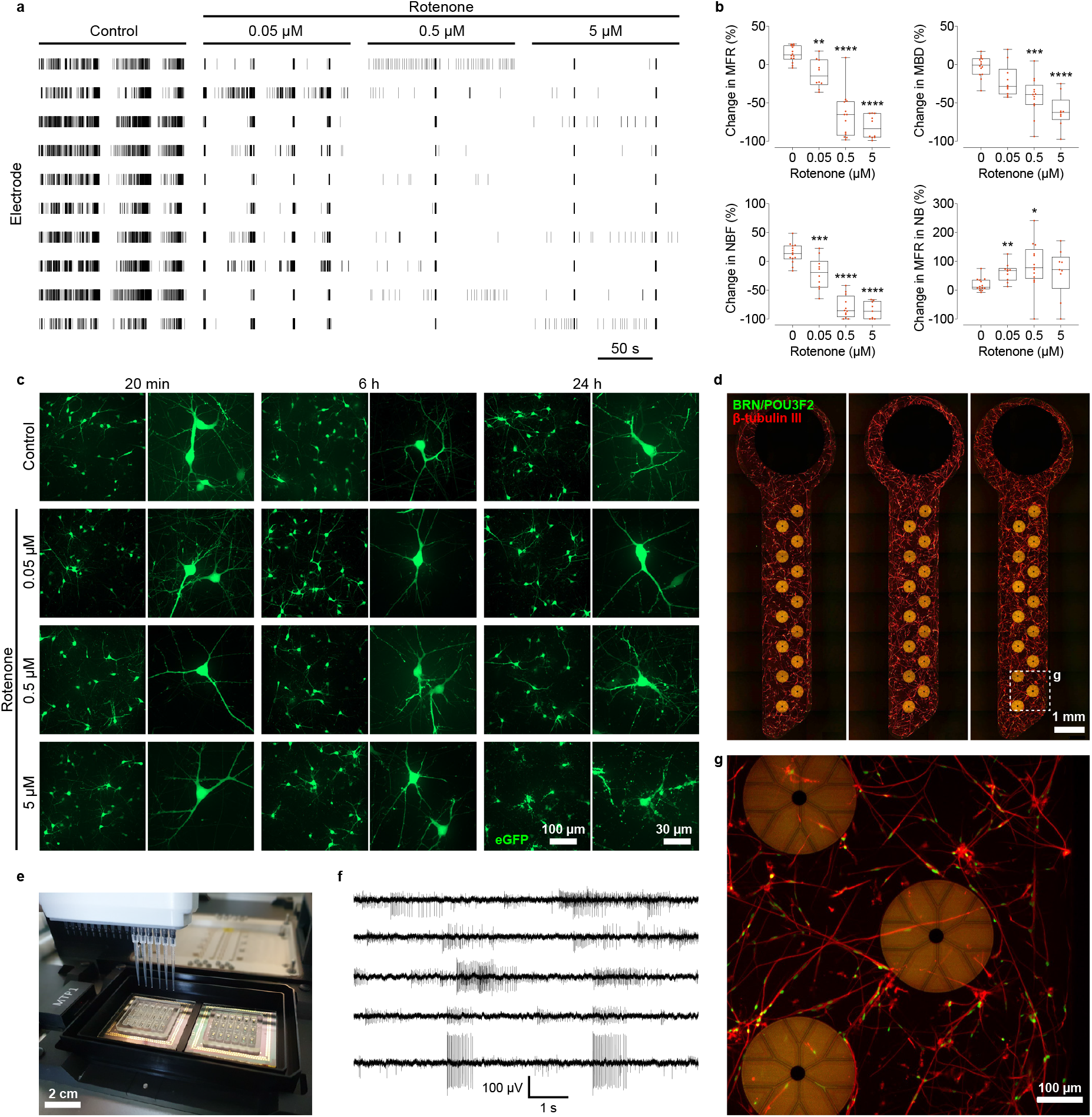
Combining morphological and functional read-outs. **a**, Raster plot of electrophysiological activity from 3D primary neurons after 10-minute treatment of rotenone at different concentrations. Black vertical lines represent one spike. For simplicity, only 10 out of 21 electrodes of one well are shown. **b**, Effect of rotenone on MFR, MBD, NBF, and MFR in NB is shown as the percentage change compared to untreated wells. The line inside each box represents the median. Whiskers indicate maximum and minimum value, and the interquartile range is set to 90%. Significance: *p<0.05; **p<0.01; ***p<0.001; ****p<0.0001. **c**, Live cell-imaging of eGFP-transduced neurons treated with rotenone for 20 min, 6 h, and 24 h. 20x images represent maximum intensity projections (MIP) of 253 to 458 µm sections; 63x images indicate MIPs of 34 to 120 µm. **d**, MIPs of smNPCs-derived dopaminergic neurons in 3 wells of the 18-well device. Cells were stained after 1 week in culture for BRN2, a key transcription factor in neuronal differentiation, and β-tubulin III. **e**, Two 18-well devices outfitted inside the 384-well microplate adapter for automatized imaging and media change. **f**, Spontaneous activity captured from smNPCs-derived dopaminergic neurons network after two weeks in vitro. Trace plots represent the spikes registered by 5 of 14 electrodes of one well in the 18-well device. **g**, MIP of 100-µm-thick section of smNPCs-derived dopaminergic neurons marked for BRN2 and β-tubulin III.

### Automating 3D in vitro screenings of human iPSC-derived neurons

Automated use of the nMPS is enabled by its microplate-compatible layout and an adapter for two nMPS for up to 36 independent experiments in parallel (Supp. video 10). Robotic systems for liquid handling (Fig. 5e) and high content imaging platforms open up the possibility to test hundreds of compounds each day while combining functional and structural data into a single platform. Automated screen was tested with small molecule–derived neural precursor cells (smNPCs) generated from commercial hiPSCs^35^, which are a powerful tool to model human neurodegenerative diseases^36^. We differentiated the smNPCs into dopaminergic neurons and cultured them in 3D following the same protocols as for primary hippocampal neurons. We confirmed that hiPSC-derived neurons also extend neurites in 3D through the tunnels, enabling the CME to record action potentials with a high SNR. Synchronized bursts demonstrated maturation of the 3D neuronal culture (Fig. 5f). After immunocytochemical labelling, a high throughput spinning disk confocal microscopy system imaged 3D neuronal cultures in all wells (Fig. 5d,g). This provides the capacity for functional and high-resolution morphological analysis of studies with large numbers of iPSC-derived neuronal lines. The multimodal nMPS described here, in combination with patient-derived iPS cells, will support a new generation of more physiologically relevant high-throughput models of neurodegenerative diseases.

## Discussion

Potential advances in treating neurological disorders are limited by our inadequate understanding of the brain’s complex structure and function. To address this gap, in vitro models are increasingly relying on 3D microphysiological cell systems. Such systems can simulate aspects of human physiology better than standard 2D approaches and reveal neuronal networks and processes at a resolution that is impossible in a whole organism. However, exploring the functional phenotype of 3D neuronal structures requires the capability to measure the electrical activity of individual neurons with millisecond resolution^37^. Commercial MEAs for 2D cultures perform poorly for 3D cultures, while recent solutions for 3D in vitro electrophysiological recordings^18,38^ are incompatible with the throughput and simple cross-platform integration required by drug discovery efforts.

Here we describe a novel technology based on substrate-integrated capped microelectrodes, allowing continuous, non-invasive electrophysiological recording from the neurites of cultured 3D neuronal cells in a re-usable, multiwell glass microfluidic device. The thin transparent glass base allows imaging of 3D neuronal structures by confocal microscopy at different time points, providing morphological details of cell–cell interactions, neurite outgrowth and synaptic elements. Likewise, live imaging reveals morphology as well as activity by calcium imaging at arbitrary locations of the 3D neuronal culture. Acquiring this information simultaneously with electrical activity recorded by CME can examine how activity patterns spread across neuronal networks. In addition, we showed how optogenetic stimulation could trigger periods of high-frequency activity in 3D neuronal circuits, using light for pacing the generation of action potentials. This could open up a new approach to studying learning and memory in vitro, using for the first time 3D neuronal circuits to explore mechanisms of synaptic plasticity. By adoption of microplate standards, most handling and readout processes can be integrated into high-throughput robotic workflows, which should enhance experimental reproducibility.

Besides its unique readout capabilities, we demonstrated how the nMPS enables 3D coculture of multiple brain cell types (neurons, microglia and astrocytes). Future work with primary innate immune cells could investigate the role of neuroinflammation in neurodegeneration^5,39^. The nMPS may increase predictivity of in vitro neurotoxicity studies, as the high sensitivity of electrophysiological recordings to early signs of neurotoxicity represents a major improvement over the morphological analysis of current 3D neuronal assays^3^. Molecular readouts, such as transcriptomics, could be implemented in future applications, as single cells can be straightforwardly retrieved from the nMPS. In combination with patient-derived iPSCs, the nMPS could enhance in vitro modelling of human disorders with reference to diagnosed disease phenotypes. We additionally envision the possibility of integrating neural organoids while using the CME to measure electrical activity. The nMPS developed here is a promising in vitro platform for disease modelling and personalized medicine, with multimodal readouts for pharmacological responses of engineered 3D cultures^4^.

## Materials and methods

### Microfabrication

Monolithic microfluidic chips were designed with Autodesk Inventor Software (Autodesk Inc. CA, USA), and fabricated in quartz by FEMTOprint SA (Switzerland) using a selective laser-induced etching process (SLE)^19,40^. MEAs and electrode caps were designed with CleWin (WieWeb, the Netherlands) and produced in house at NMI. Glass MEAs (1 mm thick) comprised transparent indium tin oxide (ITO) electrical paths insulated by silicon nitride. Microelectrodes and reference electrodes were gold.

CMEs were produced in two photolithographically structured layers of permanent epoxy photoresist (supplementary figure 1). MEAs were pretreated with oxygen plasma (2 min, 1800 mPa) then baked in an oven for 1 h at 150°C. SU-8 2002 (MicroChem, MA, USA) was spin-coated to a thickness of 3 µm (10 s at 500 rpm then 30 s at 1000 rpm). A soft bake was performed for 5 min at 95°C on a hotplate, ramped from and to room temperature. Structures were exposed (250 mJ/cm^2^, i-line, soft contact) through a photomask using a SÜSS MA6 mask aligner. A post-exposure bake (PEB), identical to the pre-exposure bake, cross-linked the exposed structures. Structures were developed in mr-Dev 600 (Micro Resist Technology GmbH, Berlin, Germany) for 15 s and then rinsed in isopropanol. An intermediate hard bake (at 150°C in an oven) further cross-linked the SU-8; this was necessary to avoid an overdevelopment of SU-8 in subsequent steps. For the second layer, ADEX TDFS A20 (Micro Resist Technology GmbH, Berlin, Germany) was laminated using a pouch laminator (GMP Photonex@325, EF02015) at 3 mm/s and at 78°C. The carrier foil was removed, and ADEX was exposed (600 mJ/cm^2^, i-line, 10 µm proximity exposure). The PEB was 1 h at 65°C, ramped. ADEX was developed in cyclohexanone for 8 min, followed by rinsing with isopropanol and drying with nitrogen.

The structure was finally hard-baked in an oven at 170°C for 30 min (ramping up and down at 0.5°C/min) to complete cross-linking for chemical stability^41^ and to prevent toxicity^42^.

The microfluidic chip was then coated with (3-aminopropyl)triethoxysilane (Sigma Aldrich) and glued to MEA with EPO-TEK 301-2FL (Epoxy Technology, Billerica, MA, USA) and aligned with a custom-made aluminum adapter. The glue was cured in an oven at 80 °C for 3 h (ramping up and down at 0.5°C/min).

### Neuronal simulations

The neural activity was simulated using LFPy 2.0^27^, built on top of the NEURON simulator^28^. The cell morphology was an interneuron from rat hippocampus, downloaded from NeuroMorpho.org (ID: NMO_00787)^43,44^. The original cell model did not have a reconstructed axon. Therefore, an axon was added manually (uniform diameter: 0.5 µm), such that it originated from the soma and passed through the tunnel. The active conductances needed to produce action potentials were inserted into the axon and soma, based on the parameters of the unmyelinated axon model from Hallermann S. *et al*.^26^. The dendrites were given the same passive parameters but no active conductances. An action potential was evoked by a single strong synaptic input to the soma compartment (synapse model: Exp2Syn, rise time constant: 0.1 ms, decay time constant: 0.1 ms, synaptic weight: 0.1 µS). The conductivity of 75% Matrigel was measured at 37°C using a conductivity measuring device (GREISINGER, GMH 3400).

The Finite Element Method (FEM) was used to make a numerical model of the recording set-up, and the general formula was similar to that used by Ness *et al*.^45^. The mesh was made with the open-source meshing tool Gmsh^46^, and the FEM simulations used FEniCS^47^. The model consisted of a large cylinder (radius: 1000 µm, height: 1000 µm) with a conductive medium (hydrogel with neurons), and the conductivity was set to σ=1.46 S/m based on conductivity measurements. The capped electrode located at the bottom center was represented as a non-conducting cylinder (radius: 125 µm, height: 25 µm), with a tunnel through the bottom (tunnel height and width: 5 µm, length: 250 µm). Using the quasi-static approximation^48^, the electric potential, φ, in a medium with conductivity, σ, can be found by solving the Poisson equation^49^, ∇·σ∇φ(r,t) =−ΣC(r,t), where ΣC(r,t) is the sum of all current sources stemming from the neuronal transmembrane currents. The recording electrodes were treated as ideal point electrodes. The simulations of the neuronal activity and the FEM simulations were done separately, and the current sources from the neuronal simulations were treated as point sources by FEniCS in the FEM simulations^45^. All outer boundaries were treated as insulating (no currents escaping the medium), corresponding to homogeneous Neumann boundary conditions. Further, the upper and side boundaries of the outer cylinder were set to ground (φ=0), corresponding to Dirichlet boundary conditions.

An analytical solution can be found for the electric potential stemming from neuronal activity in a semi-infinite conductor above an insulating plane^45^. This solution can be expected to be quite similar to the numerical FEM solution sufficiently far away from the insulating tunnel structure. Therefore, as a control, the potential simulated with FEM was compared to the analytical solution at x = −180 µm (MEA electrode location outside the tunnel), that is, 55 µm away from the tunnel structure. The maximum relative error between the FEM and analytic solution was here 5.4%.

Code to reproduce these results is available from: https://github.com/torbjone/MEA_tunnel_FEM/

### 3D neuronal culture

Dissociated 3D cell cultures were prepared from brains isolated from Swiss mice (Janvier Labs, France). All procedures were conducted in accordance with the European Union (EU) legislation for the care and use of laboratory animals (Directive 2010/63/ EU, of the European Parliament on the protection of animals used for scientific purposes, German TierSchG (Tierschutzgesetz) with latest revision 2019).

Neurons and astrocytes were isolated from embryonic hippocampi at E16-17 as previously described^50,51^. Cell dissociation was performed following the Neural Dissociation Kit (T) (Miltenyi Biotec GmbH, 130-093-231). Plating media was Neurobasal Plus Medium with 2 % (v/v) of SM1 Neuronal Supplement (STEMCELL, 5711) and 0.5 % (v/v) of Penicillin-Streptomycin (Sigma, P4333). Before plating cells, devices were plasma-treated for 2 min to sterilize and hydrophilize the surfaces. Resuspended cells were counted (NucleoCounter NC-200, Chemometec), mixed 1:3 with Matrigel® Growth Factor Reduced (Corning, 356230) and kept on ice during seeding to avoid Matrigel polymerization. Simultaneously, the devices were kept cold during plating by placing them on top of a cold rack. By using cold low retention tips, 9 µl of cell solution was dispensed in each well of the device and immediately incubated at 37 °C to promote Matrigel polymerization. The standard concentration of cells employed was 3000 per µl. After polymerization, 50 µL of medium was added to the liquid layer. 3D cultures were maintained for two weeks, adding fresh medium (1:1) every 2 days. Starting at 6 days in vitro (DIV), Neurobasal Plus Medium was replaced (1:1) by BrainPhys Neuronal Medium (STEMCELL, 05790) with 2 % (v/v) of SM1 Neuronal Supplement and 0.5 % (v/v) of Penicillin-Streptomycin.

### Glial culture preparation

Brains from postnatal Swiss mice (day 1-2) (Janvier Labs, France) were dissected, freed from meninges, minced and prepared following the protocol of Garcia-Agudo *et al*.^52^. Minced samples were digested with trypsin and EDTA 0.05% (Sigma, T4049) for 10 min at 37°C. To stop the enzymatic digestion, microglia medium containing DMEM (Gibco, 31966-021) with 10% horse serum (Gibco, 26050070) and 0.5% Penicillin-Streptomycin was added to the brains in addition to 400 IU/ brain of DNase I (Cell systems, LS002139). Samples were centrifuged (10min, 150g, RT) after mechanical trituration and plated in a Poly-D-Lysine (PDL)-coated (50 µg/ml) T75 flask. Glia culture was then incubated at 37°C and 5% CO_2_. Medium was changed after 1 and after 2 days in vitro. To promote microglial growth, cells were stimulated with L-929 conditioned medium (1:2) after 4 DIV and 7 DIV. Cells were harvested after 9 DIV by manually shaking the flask. After sorting the cells via flow cytometry, pure microglia were seeded in culture together with neurons and astrocytes (1:8).

### L-929 conditioned medium

L-929 mouse fibroblasts were seeded in 75 cm^2^ flasks at a density of 4.7×10^5^ cells and incubated in L929 medium (DMEM, 1% Penicillin-Streptomycin, 10% FBS) for 9 days at 37°C and 5% CO_2_. Medium from flasks was collected, filtered through a 0.22 µm filter and stored at −20°C until use.

### Flow cytometry

To ensure microglia purity, cells were collected from the flask, centrifuged (10 min, 150g, RT) and blocked for Fc receptors by using antibody CD16/32 (Invitrogen, 14-0161-82) (1:100). After washing the sample with washing buffer (PBS, 1% Bovine Serum Albumin (BSA), cells were centrifuged (10 min, 150g, RT) and stained for microglia marker CD11b (Invitrogen, 11-0112-81) (1:100). For dead cell exclusion a 7-AAD viability staining solution was employed (Biolegend, 420403) (5 µL per million cells). Filtered samples were acquired using the BD FACSMelody Cell Sorter (BD Biosciences) and then centrifuged for 10 min at 150 g before counting cells. Isolated microglia cells were co-cultured with isolated neural cells in Neurobasal Plus Medium with 2 % (v/v) of SM1 Neuronal Supplement, 0.5 % (v/v) of Penicillin-Streptomycin, 100 ng/ml of IL-34 (Peprotech, 200-34) and 2 ng/ml of TGF-β2 (Peprotech, 100-35B). After 6 DIV medium was replaced by BrainPhys Neuronal Medium containing 2 % (v/v) of SM1 Neuronal Supplement, 0.5 % (v/v) of Penicillin-Streptomycin, 100 ng/ml of IL-34 and 2 ng/ml of TGF-β2.

### Human iPS cell culture

GM23280 human iPS cells were purchased from Coriell Institute for Medical Research. Human iPS cells were maintained and propagated as colonies in feeder-free conditions in Essential 8 Flex Medium (Thermo Fisher Scientific, A2858501) on Matrigel hES qualified (Corning, 354277) coated plates. Colonies were passaged as aggregates every 5-7 days. Briefly, iPSC colonies were treated with Gentle Dissociation Reagent (GDR, StemCell technologies, 07174) for 7 minutes at room temperature. GDR was then replaced with E-8 flex media and colonies were gently triturated by pipetting in order to obtain a suspension of c.a. 100 μm aggregates. Small colonies were plated in freshly coated plates, and the media was refreshed every other day.

### Virus production in HEK293TF cells

HEK293FT cells were grown in OptiMEM (Life Technologies, 31985054) medium supplemented with 10% FBS (Gibco, 10499044). Cells were seeded onto (0.01 %) Poly-L-ornithine (Sigma-Aldrich, P3655) coated T75 flasks at a density of 5.5×10^6^ cells per flask. The next day, the medium was refreshed 2 hours before transfection. The transfection mixture consisted of 6 μg transgene vector (pLV_TRET_hNgn2_UBC_Blast_T2A_rtTA3), 3 μg psPax vector (Addgene #12260), 1.5 μg pMD2.G vector (Addgene #12259) and 32 μl of Fugene HD (Promega, E2312) in 600 µl of serum-free OptiMEM (Thermo Fisher Scientific, 31985054) medium. 24 hours after transfection, the old medium was replaced with fresh medium containing 1 % FBS and 10 mM HEPES (Thermo Fisher Scientific, 15630056) (low-FBS medium). Three flasks were used for the batch of virus production. The viral supernatant was collected at 48 hours and 72 hours and centrifuged at 1000 g for 10 minutes at 4 °C to pellet cell debris. Viral supernatant was transferred into a new sterile 50 mL tube and passed through a low protein binding 0.45 μm filter (SE1M003M00, Millipore) to exclude any cells still present in the supernatant. The supernatant was then transferred into Amicon spin filter with a 100 kDa cutoff (UFC910024, Millipore) and centrifuged at 4000 g for 20 to 30 min; this step was repeated several times until all the media had been spun to yield 1 ml of concentrated virus that was stored in single-use aliquots at −80 °C.

The pLV_TRET_hNgn2_UBC_Blast_T2A_rtTA3 plasmid was engineered by combining hNgn2_UBC (from Addgene Plasmid # 61474) with rTTA3 (from Addgene Plasmid # 61472) to yield a single vector system with blasticidin resistance.

### Generation of stable smNPCs for inducible expression of Neurogenin-2

The smNPCs were derived from iPS cells using the method described in Reinhardt *et al*.^53^, applying few adjustments ^35^. The smNPCs were cultured on Matrigel (Corning, 354234) coated plates in the expansion medium (N2B27) supplemented with 3 µM CHIR (R&D System, 4423), 0.5 µM PMA (Cayman Chemical, 10009634) and 64mg/l Ascorbic acid (AA, Sigma-Aldrich, A8960) for up to 10 passages before use in any downstream experiments. N2B27 medium consisted of DMEM/F-12-GlutaMAX (Thermo Fisher Scientific, 31331093) and Neurobasal (Thermo Fisher Scientific, 21103049) mixed at 1:1 ratio, supplemented with 1:200 N-2 supplements (Thermo Fisher Scientific, 17502048), 1:100 B-27 supplements without vitamin A (Thermo Fisher Scientific, 12587010), 1:200 GlutaMAX™ (Thermo Fisher Scientific, 35050), 5 μg/ml insulin (Sigma-Aldrich, I9278), 1:200 non-essential amino acids (NEAA, Thermo Fisher Scientific, 11140050), 55 μM 2-mercaptoethanol (Thermo Fisher Scientific, 21985023), and 1:100 penicillin/streptomycin. The smNPCs were routinely passaged using Accutase (Sigma-Aldrich, A6964). The smNPCs at passage 10 were transduced with pLV_TRET_hNgn2_UBC_Blast_T2A_rtTA3 viral particles to produce the doxycycline-inducible Neurogenin-2 stable cell line. A multiplicity of infection (MOI) of 60 was used for the viral transduction. The cells were subsequently selected with blasticidin (15 µg/ml, InvivoGen, ant-bl-1) for 3 passages or up to 15 days.

### smNPC differentiation

NGN2-transduced smNPCs were expanded for 5 to 7 days before differentiation in smNPCs expansion medium on Matrigel-coated plates. After reaching 70-80 % of confluency, smNPCs colonies were dissociated by Accutase treatment for 20 min at 37 °C, collected, and washed in N2B27 medium with centrifugation at 300 g for 3 min at room temperature. Cells were then resuspended in N2B27 medium supplemented with 2.5 µg/ml doxycycline (Dox, Sigma-Aldrich, D9891), 3 µM CHIR, 64 mg/l AA, 0.5 µM SAG (Merck Millipore, 566660). Cells were counted with trypan blue in the Biorad – TC-20 automatic cell counter and seeded at 850k/well in N2B27 medium with 2.5 µg/ml Dox, 3 µM CHIR, 64 mg/l AA, 0.5 µM SAG onto Matrigel-coated 6-well plates. Cells were left in the incubator overnight to promote adhesion. Two days after seeding, medium was replaced with N2B27 medium supplemented with 2.5 µg/ml Dox, 0.7 µM CHIR, 64 mg/l AA, 0.5 µM SAG, 100 ng/ml FGF8 (Peprotech, 100-25). Four to five days after seeding, cells were dissociated in Accutase at 37°C for 20min, collected and washed in N2B27 medium (w/o supplements) and centrifuged at 300 g for 3 min. Cells were resuspended in 500-600 μl of N2B27 medium containing 10 μM DAPT, 64 mg/l AA, 10 ng/ml BDNF (PeproTech, 450-02), 10 ng/ml GDNF (PeproTech, 450-10), 500 µM cAMP (AppliChem, A0455), 2.5 ng/ml Activin A (PeproTech, AF-120-14E), 1 ng/ml TGF-b3 (PeproTech, 10036E), 0.5 µg/ml laminin (Sigma-Aldrich, L2020). Viable cells were counted and diluted 1:4 in Matrigel reduced growth factor (Corning, 356230) to achieve a cell density of 2000 to 6000/µL. The cell suspension was plated in 3D in the nMPS devices as described above. Half media change was performed every 2-3 days, and cells were in culture up to two weeks after seeding in 3D.

### AAV1-eGFP transduction

One day after plating, neurons growing in 3D were transduced with adeno-associated viruses serotype 1 (AAV1) carrying a transgene for the expression of enhanced green fluorescent protein (eGFP) under the human Synapsin 1 (hSyn) promoter. pAAV-hSyn-EGFP was a gift from Bryan Roth (Addgene catalog #50465-AAV1). The MOI utilized was 10^5^, and the expression of eGFP was monitored daily via microscopy. After 4 DIV, 3D cultures were homogeneously labeled in green with no apparent negative effect on viability.

### Immunocytochemistry

3D neuronal cultures were fixed with 4 % (w/v) PFA and 4 % (w/v) glucose solution at 37 °C for 20 min and rinsed five times with washing buffer (PBS −/−with 0.2% fish skin gelatin (G7765, Sigma)). Afterwards, the 3D cultures were permeabilized with PBS (−/−) supplemented with 0.3% (w/v) Triton X-100 and tilted on a rocker at RT for 1 h. The permeabilization buffer was rinsed five times with washing buffer, replaced with blocking buffer (PBS (−/−) with 0.2 % (v/v) donkey serum and 0.1% (w/v) Triton X-100), and incubated on a rocker overnight at 4 °C. The blocking buffer was discarded and a primary antibody solution was added, incubating samples on a rocker overnight at 4 °C. The primary antibody was then washed off overnight on a rocker at 4 °C, followed by incubation with the secondary antibody on a rocker overnight at 4 °C. Finally, the secondary antibody was rinsed off, adding mounting medium Slowfade Diamond (2054439, Thermo Fisher Scientific, MA, US) to improve the fluorescence retention, preservation over time and to label the nuclei with DAPI.

Primary 3D cultures were stained for GFAP (mouse, anti-GFAP, 173 011, Synaptic Systems), Homer-1 (mouse, Anti-Homer 1, 160 003, Synaptic Systems), Iba-1 (mouse, anti-Iba1, 019-19741, FUJIFILM Wako Chemicals), MAP2 (mouse, anti-MAP2, NB300-213, Novus Biologicals), SMI-1 (mouse, anti-SMI132, 837904, BioLegend) and β-tubulin III (mouse, anti-β-tubulin III, 801201, BioLegend).

smNPC-neurons were stained for β-tubulin III (anti-β-tubulin III, T2200, Sigma Aldrich) and BRN2 (anti-BRN/POU3F2, 12137, Cell Signaling) after 1 week in culture.

### Imaging of 3D cultures

Imaging and morphological analysis of 3D primary cultures were performed after one week in vitro. High-resolution images were taken with a Zeiss LSM 780 NLO confocal equipped with an Airyscan unit microscope stand. For imaging large areas of the hydrogel, a Zeiss Cell Observer® System with spinning disk head was employed. The software used for capturing the images was ZEN (blue edition, ZEISS). After the acquisition, images were processed and reconstructed in IMARIS 4.6 (Bitplane AG, Oxford Instruments).

For automated imaging, a customized 384-well microplate adaptor was designed. This platform was produced by Weerg Srl (Italy) and supports the analysis of two devices at once. The robotic system contains a Microlab Star that performs all liquid handling and an integrated Cytomat-24 CO_2_ incubator^35^. The plates were transported within the system by the robotic arm. High-content screening of stained smNPC-neurons was performed with a high content automated spinning disk confocal microscope (Yokogawa, CV7000). Images were then processed and exported using the Cell PathFinder software (Yokogawa, version 3.03.02.02).

### Electrophysiological recordings

The electrical activity of the 3D neuronal network was measured by a USB-MEA 256-System (Multi Channel Systems MCS GmbH, Germany), allowing simultaneous recording of 256 channels at a maximum sampling rate of 50 kHz per channel. The system has an integrated filter amplifier with gain and bandwidth adjustable in the acquisition software. Recording of spontaneous activity of neuronal circuits was performed after ten days in vitro.

Experiments were carried out whilst continuously monitoring temperature, CO_2_, and humidity using a customized MEA incubation chamber (Okolab Srl, Italy). Recordings started 10 min after closing the chamber to allow stabilization of environmental conditions. The temperature was set at 36 °C, the humidity at 85 %, and CO_2_ at 5 %.

Spike detection, burst, and network burst analysis were performed using NeuroExplorer (version 5.300). First, recordings were filtered with a fourth-order bandpass filter (60-6000 Hz). Then, action potentials were detected with a threshold set at T = 4σ_N_, where σ_N_ is an estimate of the standard deviation of the background noise^54^. The minimum time between spikes was set to 1 ms. Burst detection was performed using a MaxInterval method. This computational method outperformed other algorithms previously used and was selected for its suitable properties as a burst detector^55^.

Within this algorithm, the minimum duration of a burst was set to 10 ms, and the lowest number of spikes that conform a burst to 5. Also, the minimum time between two signals from two individual bursts was fixed to 200 ms and the maximum time between spikes in a burst to 170 ms at the beginning of the burst and 300 ms within the burst. These values were optimized via the analysis of multiple recordings of 2D and 3D neuronal cultures.

Network bursts within each well of the MPS device were identified by a customized script (Python). This script calculated the median of each burst in every electrode and projected them as single events in one timeline. Network bursts were then defined by Poisson distribution of the medians with a surprise value of 3 and a minimum duration of 5 ms. The duration of the network bursts was subsequently calculated by averaging the first and the last spikes of each burst that was part of the network burst. The minimum number of electrodes contributing to the network burst was fixed to 33.3 % of the total electrodes, i.e. 6 electrodes per well.

### Ca^2+^ imaging

Neuronal cultures were transduced 1-3 days after plating with an adeno-associated virus expressing GCaMP6f as Ca^2+^ indicator. pAAV.Syn.GCaMP6f.WPRE.SV40 was a gift from Kim Douglas (Addgene catalog #100837-AAV1). A week later, the samples were placed inside the MEA incubation chamber and illuminated with a broad-spectrum LED illumination system (CoolLED pE-300ultra, CoolLED, UK). Fluorescence images were acquired at 5 full frames per second (50 ms acquisition time) by an iXon EMCCD camera (Oxford Instruments) mounted on an inverted microscope (ECLIPSE Ti2, Nikon GmbH). Different sections of the 900 μm thick hydrogel were imaged for 2-3 min with a 20x objective. The electrical activity was simultaneously recorded by the MEA system at 50 kHz.

The total fluorescence of each frame, proportional to intracellular Ca^2+^ concentration, was expressed in arbitrary units, which were calculated using NIS-Elements software (Nikon Instruments Inc.). The timing of imaging experiments could be linked to acquisition of electrophysiological data by internal triggering via the recording software.

### Optical stimulation

Channelrhodopsin-2 (hChR2) was expressed in 3D neuronal cultures by transduction with an adeno-associated virus. pAAV-hSyn-hChR2(H134R)-EYFP was a gift from Karl Deisseroth (Addgene catalog #26973-AAV1). At ten days in vitro, the 3D neuronal network was illuminated with blue light (470 nm) to activate hChR2. Simultaneously, the evoked electrical activity was recorded by the MEA system. The illuminated area was defined in the XY plane by a digital micromirror device system (Mosaic3, Oxford Instruments) attached to the microscope. The duration of the light stimuli was 5 ms with a frequency of 40 Hz, while the illuminated region comprised an area of 0.04 mm^2^. Optogenetic experiments were performed inside the MEA incubation chamber. Electrophysiological data acquired before, during and after light stimulation were post-processed for spike detection as described above.

### Compound application

Responses to PTX (P1675, Sigma-Aldrich), TTX (T-550, Alomone Labs), and Rotenone (R8875, Sigma-Aldrich) were monitored after adding each compound to individual wells of the device placed inside the MEA recording system. Ten percent of the culture medium was replaced by fresh medium containing test compounds 10x the final concentration and carefully mixed by an electronic multi-pipette (VIAFLO 12 channels, 12 μl, INTEGRA Bioscience). All experiments included negative controls to assess the effect of the application procedure. The response to the neuroactive compounds was recorded starting 10 min after application.

### Reusability of nMPS

Glass devices were reused several times after thoroughly washing them. They were rinsed with water and incubated for at least 3 h in bi-distilled water with Tergazyme 1 % at room temperature, followed by an overnight rinse in bi-distilled water. If necessary, the monolithic glass piece could be detached from the MEA substrate by incubation in concentrated sulfuric acid overnight. Then, it could be glued again to another MEA substrate.

### Statistical analysis

Data were analyzed using GraphPad Prism 8 (GraphPad Software Inc. v 8.3.4). Quantile-quantile plots, frequency histograms, and Shapiro-Wilks tests were employed to gauge the degree of normality of the data distribution both visually and qualitatively. Variability analysis was performed at electrode, well, device, and preparation level. Compound effects were compared to the non-treated wells. First, the variances of each group were tested for statistical differences using Bartlett’s test. If variances of groups were significantly different from each other, Brown test was employed. Then, post hoc tests Dunnett T3 (if the sample size was smaller than 50) or Games-Howell (if the sample size was bigger than 50) were applied. If variances did not show significant differences among groups, an ordinary one-way ANOVA was run. Subsequently, post hoc tests Tukey or Dunnett were used. The latter was employed if groups were compared to a control group. Data were presented either in violin-plots illustrating the median values ± quartiles or shown in box plots representing the median, and the maximum and minimum values highlighted with whiskers. Asterisks denote statistical significance as follows: *p<0.05; **p<0.01; ***p<0.001; ****p<0.0001.

## Supplementary materials

**Supplementary video 1: 3D eGFP-transduced neuron**. Live imaging of hippocampal neuron with 3D neurite outgrowth and arborization. An eGFP-expressing neuron embedded in Matrigel was imaged with a Cell Observer confocal microscope after 10 DIV. Dimensions 134×151×45 μm.

**Supplementary video 2: 3D eGFP-transduced neurite with dendritic spines**. High-resolution live image of a single dendrite expressing eGFP. Image acquired by a confocal microscope with Airyscan unit (Zeiss). In the 3D reconstruction, it is possible to visualize several spines protruding from a single dendrite.

**Supplementary video 3: Immunofluorescence staining of a 3D co-culture**. Primary neurons co-cultured with astrocytes and microglia growing in 3D were immunolabeled for nucleus-related marker DAPI (blue), IBA1 (red), GFAP (yellow), and neurite related marker β-tubulin III (green). Dimensions 112×112×109 μm.

**Supplementary video 4: Immunofluorescence staining of 3D neuronal network**. The complex 3D neurite network was imaged after the immunolabeling of axon-related marker SMI (red) and dendrite-related marker MAP2 (green). Image acquired by LSM 780 confocal with Airyscan unit on AxioObserver Z.1 microscope. Dimensions 112×112×73 μm.

**Supplementary video 5: Immunofluorescence localization of postsynaptic densities in a 3D cell culture**. Neurons growing in 3D were immunolabeled for postsynaptic density-related marker Homer 1 (blue) and neuronal marker β-tubulin III (green). Image acquired by LSM 780 confocal with Airyscan unit on AxioObserver Z.1 microscope. Dimensions 47×47×11 μm.

**Supplementary video 6: Simulation of an action potential captured by a CME**. Representation of an action potential evoked in a neuronal soma growing in 3D and transmitted through an axon trapped in one tunnel of the CME. Color indicates the extracellular potential (logarithmic scale with positive in red and negative in blue). The extracellular potential at a capped microelectrode (CME) is amplified by two orders of magnitude in comparison to the magnitude generated by an axon near an open microelectrode (OME). For comparison, the membrane potential at the soma (purple) and along the proximal (green) and distal (pink) axon. The inset graph shows the peak extracellular potential generated near the soma. A magnitude above 10 µV is reached only within a distance of 15 µm.

**Supplementary video 7: Spontaneous intracellular calcium oscillations in GCaMP6f-transduced neurons growing in the nMPS**. Synchronized oscillations of primary untreated neurons expressing GCaMP6f were captured by calcium imaging after 10 DIV. Images of 200×200 µm were acquired at 5 frames per second and at 560 µm from the bottom of the device.

**Supplementary video 8: Intracellular calcium oscillations in GCaMP6f-transduced neurons treated with PTX**. Large synchronous oscillations in intracellular calcium observed in 3D neuronal networks following application of PTX (10 µM). Images were taken at 5 frames per second, 10 minutes after applying PTX. The region of the nMPS imaged had an area of 200×200 µm and was at 560 µm from the bottom of the device.

**Supplementary video 9: Intracellular calcium oscillations in GCaMP6f-transduced neurons treated with TTX**. Intracellular calcium oscillations in 3D neuronal networks were almost completely abolished by application of TTX (10 nM). Images were taken at 5 frames per second, 10 minutes after applying TTX. The region of the nMPS imaged had an area of 200×200 µm and was at 560 µm from the bottom of the device.

**Supplementary video 10: Automated handling of nMPS**. Media change performed at the liquid handling station of the automated culture system. The pipetting arm transports three 300 µl tips in close proximity to the wells of one of the two 18-wells nMPS placed inside the custom-designed adapter to perform media aspiration and discarding it in the waste collection module. This step is repeated sequentially for all wells of each device. Fresh media is then dispensed sequentially in all the wells using a new set of 300 µl tips. Once media change is performed for all the wells of the nMPS, the robotic arm re-lids the plate and transports it back to the CO_2_ incubator. The nMPS-microplate is then transported from the liquid handling station shelf to the Yokogawa high-content imaging station using a robotic arm.

**Supplementary figure 1: Representation of the CME fabrication process on top of an MEA**. First, SU-8 was spin-coated to produce open micro tunnel structures around the electrodes (ii). SU-8 was then exposed (pink arrows) through a photomask (iii). A post-exposure bake (PEB) further cross-linked and stabilized the exposed structures. SU-8 was then developed and baked (iv). In order to confine the space over the electrodes, ADEX was laminated over the SU-8 (v) and then exposed through a photomask (vi). This was followed by a post-exposure bake. ADEX was then developed and baked (vii).

**Supplementary figure 2: Response to PTX, TTX, and rotenone**. Boxplots of percentage change of burst frequency rate (BFR), mean firing rate in bursts (MFR in B), interspike interval coefficient of variance (ISI CoV), and bursts in NB (B in NB) after applying different doses of PTX (a), TTX (b), and rotenone (c). Orange dots denote the mean value of wells to which certain concentration was applied. Whiskers illustrate the minimum and maximum value. Interquartile range is set to 90%. Asterisks indicate significant differences between the non-treated wells vs the treated wells: *p<0.05; **p<0.01; ***p<0.001; ****p<0.0001.

**Supplementary figure 3: Electrical activity recorded in single wells from three different devices**. Values of each electrode were grouped by wells and compared to the mean of all the values of a 12-well device (A, C or D). All data were obtained from one single preparation and collected at 10 DIV. The following parameters are represented in violin plots: mean firing rate (MFR) (Hz), burst frequency rate (BFR) (Hz), number of spikes in burst (SiB) (%), burst duration (BD) (s), and interspike interval of the coefficient of variance (ISI CoV). Values of single CME are shown as red dots, and the mean of all values from a certain device is indicated by a discontinuous black line. Median and quartiles are illustrated inside the violin plots as a continuous black line. Asterisks indicate significant differences between a well with the mean of all values of a certain device: *p<0.05; **p<0.01; ***p<0.001; ****p<0.0001.

**Supplementary figure 4: Distribution of the electrophysiological parameters**. Collected data (N = 187 wells) of spontaneous activity recorded under control conditions in different devices is represented in QQ plots and frequency histograms. In the QQ plots, the red discontinued line indicates a perfect fit for the data distribution. The following data are represented: mean firing rate (MFR) (Hz), burst frequency rate (BFR) (Hz), number of spikes in burst (SiB) (%), burst duration (BD) (s), interspike interval of the coefficient of variance (ISI CoV), network burst frequency (NBF) (Hz), mean firing rate in network bursts (MFR in NB) (Hz) and number of bursts in network bursts (B in NB) (%).

**Supplementary figure 5: Data represented according to preparations**. Means of all wells were categorized into the different days of preparation and analyzed for significant differences with the mean of all values. The following data are represented in violin plots: mean firing rate (MFR) (Hz), burst frequency rate (BFR) (Hz), number of spikes in burst (SiB) (%), burst duration (BD) (s), interspike interval of the coefficient of variance (ISI CoV), network burst frequency (NBF) (Hz), mean firing rate in network bursts (MFR in NB) (Hz) and number of bursts in network bursts (B in NB) (%). Means of wells are denoted by orange dots. The mean of all values is indicated by a discontinuous black line. Median and quartiles are illustrated inside the violin plots as a continuous black line. Asterisks indicate significant differences between a certain day of preparation and the mean of all values: *p<0.05; **p<0.01; ***p<0.001; ****p<0.0001.

**Supplementary figure 6: Data represented according to used devices**. Means of all wells were categorized into the 5 different 12-well devices used for this study and analyzed for significant differences with the mean of all values. The following data are represented in violin plots: mean firing rate (MFR) (Hz), burst frequency rate (BFR) (Hz), number of spikes in burst (SiB) (%), burst duration (BD) (s), interspike interval of the coefficient of variance (ISI CoV), network burst frequency (NBF) (Hz), mean firing rate in network bursts (MFR in NB) (Hz) and number of bursts in network bursts (B in NB) (%). Means of wells are shown as green dots. The mean of all values is represented as a discontinuous black line. Median and quartiles are illustrated inside the violin plots as a continuous black line. Asterisks indicate significant differences between a certain device and the mean of all values: *p<0.05; **p<0.01; ***p<0.001; ****p<0.0001.

**Supplementary figure 7: Data of single devices categorized according to preparations**. Means of wells of four devices (A, B, C and D) corresponding to three different preparations (1, 3 and 4) were compared to the mean of all the values of a given preparation. The following data are represented in violin plots: mean firing rate (MFR) (Hz), burst frequency rate (BFR) (Hz), number of spikes in burst (SiB) (%), burst duration (BD) (s), interspike interval of the coefficient of variance (ISI CoV), network burst frequency (NBF) (Hz), mean firing rate in network bursts (MFR in NB) (Hz) and number of bursts in network bursts (B in NB) (%). Means of wells are denoted by yellow dots. The mean of values from a certain day of preparation is indicated by a discontinuous black line. Median and quartiles are illustrated inside the violin plots as a continuous black line. Asterisks indicate significant differences between a certain device of a certain day of preparation and the mean of all values collected that day: *p<0.05; **p<0.01; ***p<0.001; ****p<0.0001.

**Supplementary table 1: Comparison of the two nMPS developed in this study**. This table summarizes the architecture and microfluidic dimensions of the 12-well and the 18-well nMPS.

## Author contributions

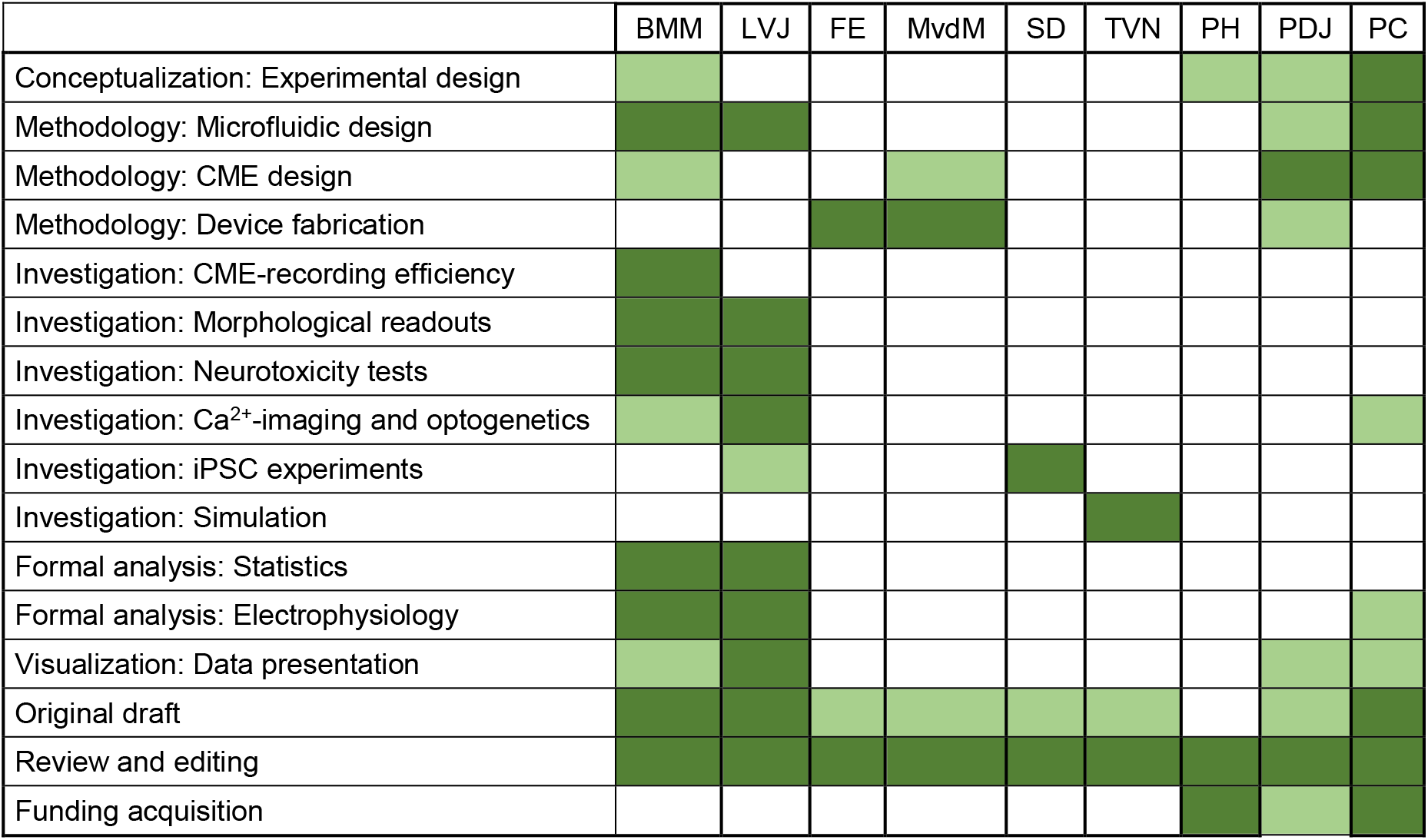

## Competing interests

BMM, PDJ and PC are named as inventors on patent applications EP3494877 and WO2019115320 (“Device for the examination of neurons”) filed by NMI Natural and Medical Sciences Institute at the University of Tübingen.

## Acknowledgments

PC and PDJ acknowledge funding from the State Ministry of Baden-Württemberg for Economic Affairs, Labour and Housing Construction. PC acknowledges funding from the German Ministry of Education and Research (BMBF) under grant agreement 031L0061 (MEAFLUIT). PC and PH acknowledge funding from Baden-Württemberg Stiftung GmbH under grant agreement MIVT-7 (NEWRON-3D). PH acknowledges funding from the Network of Centres of Excellence in Neurodegeneration (CoEN), under grant agreement 5010. TVN acknowledges funding from the European Union Horizon 2020 Framework Programme for Research and Innovation under Specific Grant Agreement No. 945539 (Human Brain Project; SGA3). MvdM acknowledges funding from the European Union Horizon 2020 Framework Programme for Research and Innovation under the Marie Sklodowska-Curie Grant Agreement No. 814244.

We thank Andrea Lovera and Rosanna Toscano (FEMTOprint, Muzzano, CH) for help with glass microfluidics, Christian Feldhaus, Vanessa Carlos and Aurora Panzera (Max Planck Institute for Developmental Biology, Tübingen, DE) for help with confocal microscopy, Anita Niedworok, Kathrin Stadelmann, Matthew McDonald, Han Liu, Angelika Stumpf and Gerhard Heusel (NMI) for technical assistance. We also thank Ashutosh Dhingra (DZNE) for sharing protocols for generation of smNPCs, Joachim Taeger (DZNE) for the set up of the automated culture, Laura Fernández García-Agudo (Max Planck Institute for Experimental Medicine, Göttingen, Germany) for help with preparation of mouse microglia, and Alexander Kirillov (NeuroExplorer, Colorado Springs, USA) for help with spike data analysis.

## Notes

https://github.com/torbjone/MEA_tunnel_FEM/

### Competing Interest Statement

Beatriz Molina-Martinez, Peter D. Jones and Paolo Cesare are named as inventors on patent applications EP3494877 and WO2019115320 (Device for the examination of neurons) filed by NMI Natural and Medical Sciences Institute at the University of Tuebingen.

