## Supplementary figures and images for "A multimodal 3D neuro-microphysiological system with neurite-trapping microelectrodes"

### Supplementary Figures 1-7

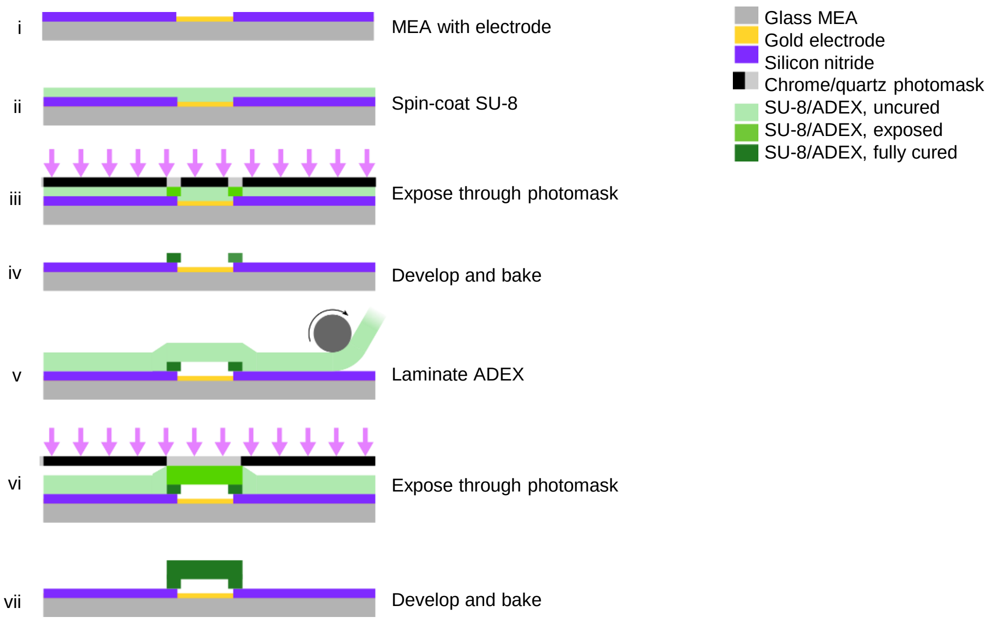

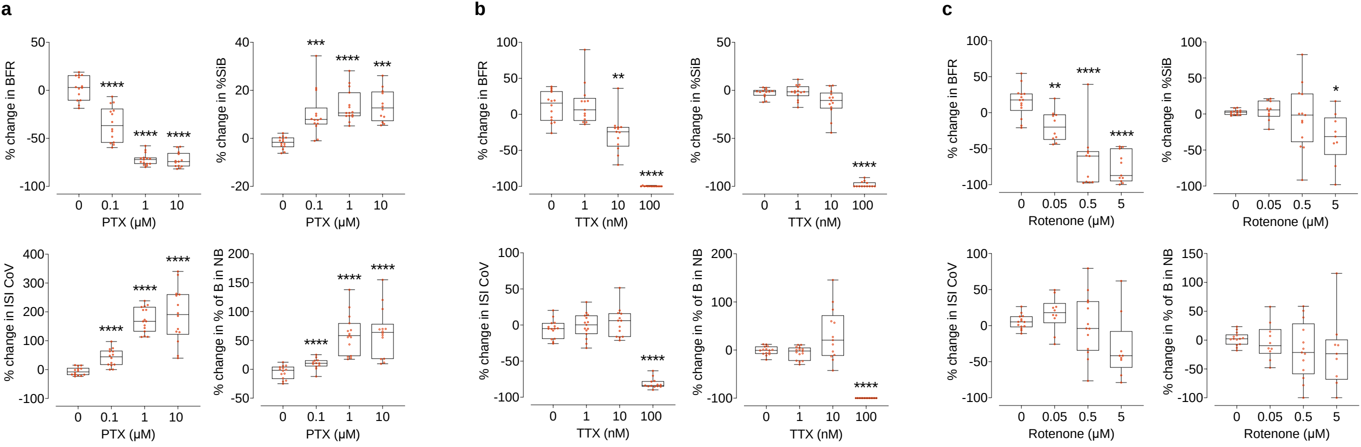

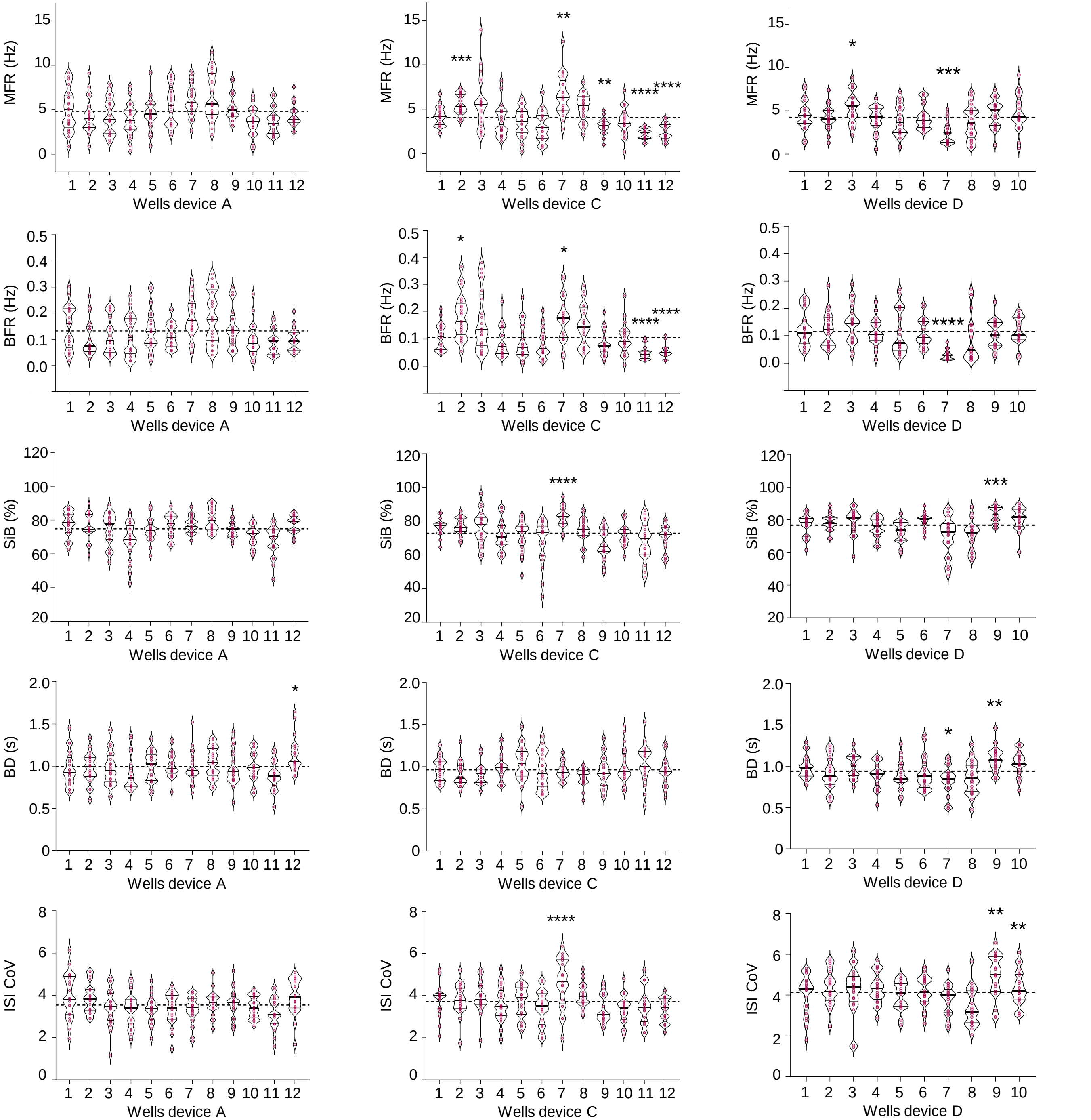

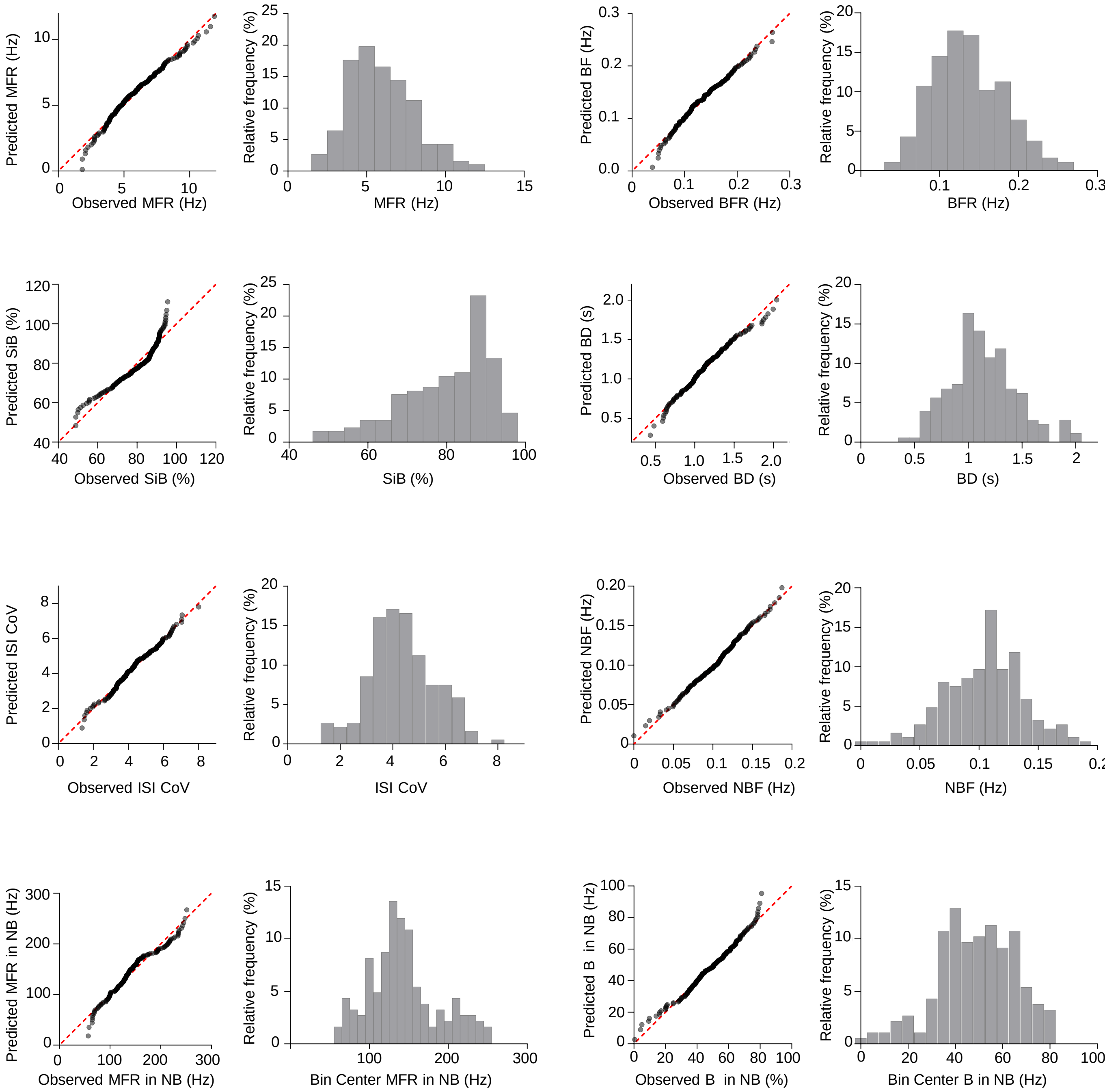

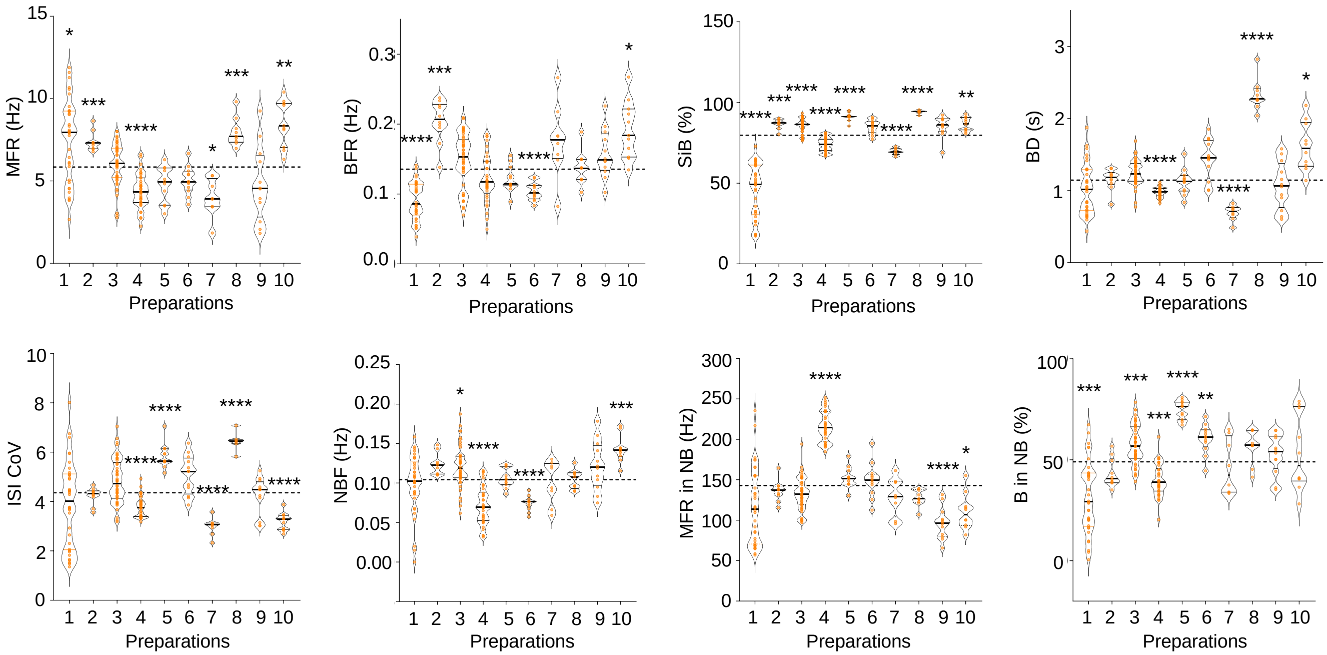

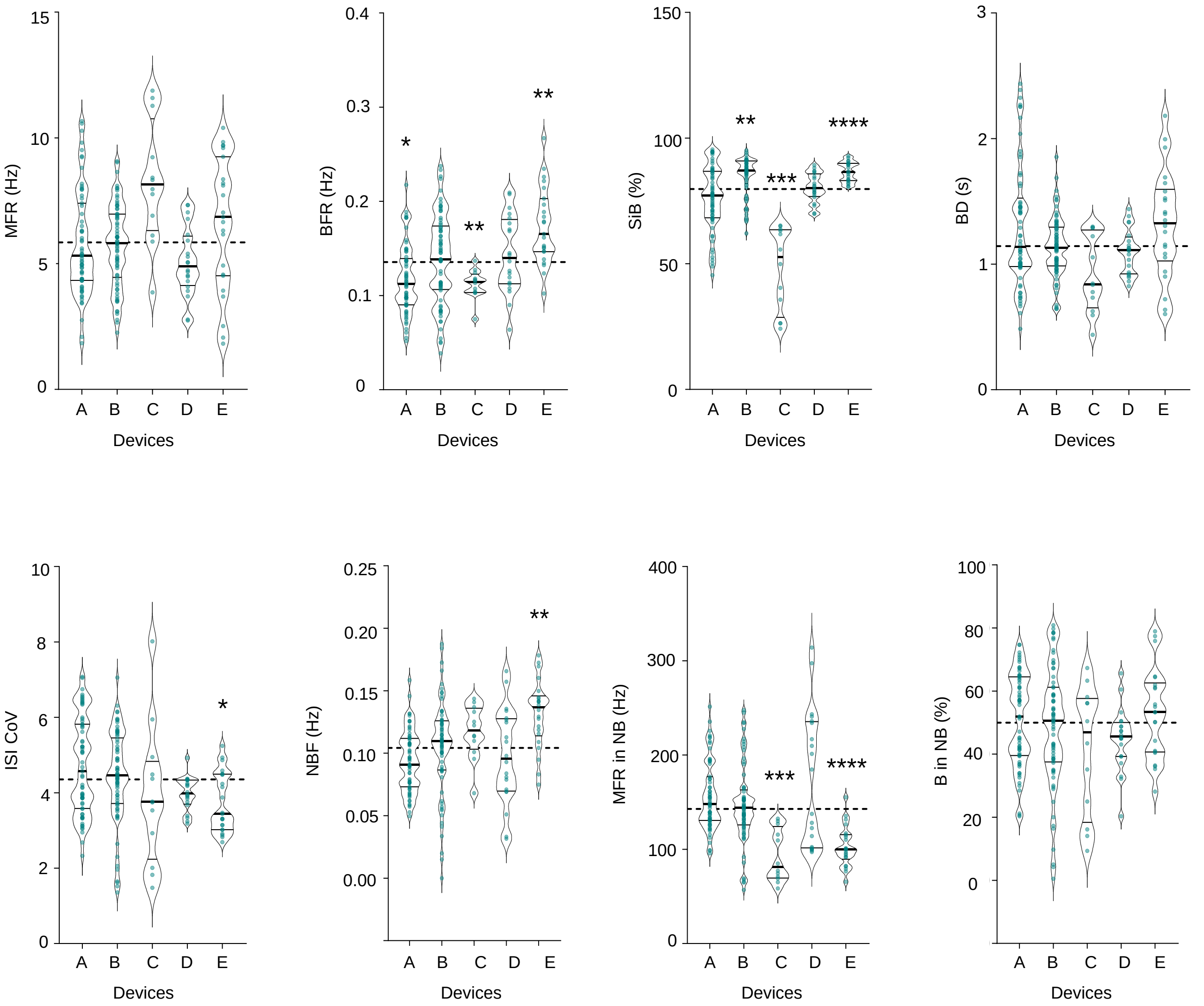

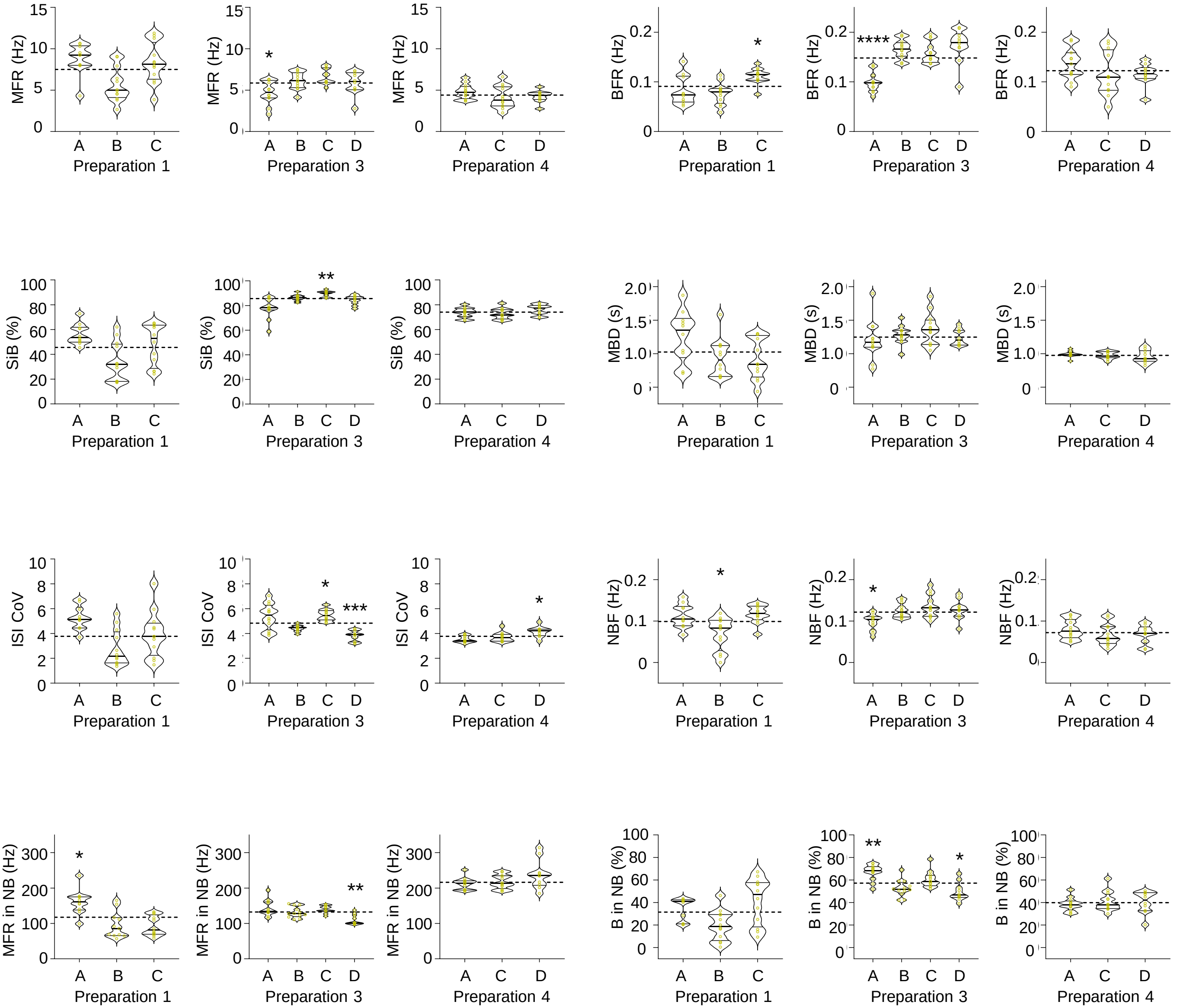
