## Supplementary Table 1 for "A multimodal 3D neuro-microphysiological system with neurite-trapping microelectrodes"

| | | 12 – well device | 18 – well device |
|---|---|---|---|
| | Number of electrodes in each well | 21 | 14 |
| | Number of microtunnel entrances | 24 | 12 |
| **Liquid layer** | Height (mm) | 2.1 | 2.1 |
| | Width (mm) | 3.5 | 3.5 |
| | Length (mm) | 12.25 | 7.8 |
| | Volume (µl) | 50 | 50 |
| **Gel layer** | Height (mm) | 0.9 | 0.9 |
| | Width (mm) | 1 | 1 |
| | Length (mm) | 8 | 5.5 |
| | Volume (µl) | 9 | 7 |
